# Identifying regulatory and spatial genomic architectural elements using cell type independent machine and deep learning models

**DOI:** 10.1101/2020.04.19.049585

**Authors:** Laura D. Martens, Oisín Faust, Liviu Pirvan, Dóra Bihary, Shamith A. Samarajiwa

**Affiliations:** MRC Cancer Unit, Hutchison-MRC Research Centre, Cambridge Biomedical Campus, University of Cambridge, Cambridge, CB2 0XZ, UK

## Abstract

Chromosome conformation capture methods such as Hi-C enables mapping of genome-wide chromatin interactions and is a promising technology to understand the role of spatial chromatin organisation in gene regulation. However, the generation and analysis of these data sets at high resolutions remain technically challenging and costly. We developed a machine and deep learning approach to predict functionally important, highly interacting chromatin regions (HICR) and topologically associated domain (TAD) boundaries independent of Hi-C data in both normal physiological states and pathological conditions such as cancer. This approach utilises gradient boosted trees and convolutional neural networks trained on both Hi-C and histone modification epigenomic data from three different cell types. Given only epigenomic modification data these models are able to predict chromatin interactions and TAD boundaries with high accuracy. We demonstrate that our models are transferable across cell types, indicating that combinatorial histone mark signatures may be universal predictors for highly interacting chromatin regions and spatial chromatin architecture elements.

## Introduction

Interactions between cis-acting regulatory elements and trans-acting regulatory proteins are facilitated by spatial genome organisation, chromatin accessibility, epigenetic modifications and non-coding RNA interactions that contribute to the complexity of gene regulatory programmes. Our genetic material, in the form of nuclear DNA, is organised into chromosomes, with each DNA molecule wrapped up and compressed into super-coiled material known as chromatin. Various studies have demonstrated that this packaging is not merely useful to compact the DNA but also plays an important role in gene regulation [1, 2, 3]. At the simplest level, DNA is wrapped around nucleosomes forming a “beads on a string” structure. These nucleosomes are made of two copies each of H2A, H2B, H3 and H4 histone proteins with 146 nucleotides of DNA wrapped around each nucleosome in 1.67 superhelical left handed turns [4]. Each nucleosome is connected to each other by short linker DNA associated with linker histone H1. The N-terminal tails of these histone proteins can be post-translationally modified, with more than 80 modifications (“histone marks”) identified so far [5, 6, 7]. These modifications act in a yet to be deciphered combinatorial code. However, the association of some of these modifications in forming active or inactive states as well as condensed heterochromatin or accessible euchromatin are known [8].

Chromatin forms both global [9] and local three–dimensional architectures; an example being the looping of DNA to bring distal regulatory regions such as enhancer and promoter into close spatial proximity [10, 11, 12, 13, 14]. More recently, chromosome conformation capture (3C) methods such as 4C [15], 5C [16] and Hi-C [17] as well as immunoprecipitation based ChIA-PET [18] have provided insight into genome scale chromatin interactions and spatial genomic organisation [19]. Analysis of Hi-C data revealed the general hierarchical organisation of chromatin, which includes structures such as A/B compartments [20], topologically associated domains (TADs) [21, 22], and long-distance chromatin loops [23].

TADs are rosette–like nested structures formed by localised chromatin interaction regions in the 0.2-1.0 Mb range. Their main characteristic is the preferential interactions of TAD enclosed chromatin regions with each other. This feature is distinctly recognisable in Hi-C contact maps as patterns of interacting loci along the diagonal. It has been proposed that TADs have an insulating function that facilitates inter–domain interaction of regulatory elements while restricting most of the cross–boundary enhancer promoter interactions (EPIs) [22, 24]. However, more recent studies show that some EPIs can form intra–TAD interactions by escaping TAD boundary insulation [25, 26]. Evidence for the role of TADs in gene regulation is seen in both normal physiological as well as pathogenic situations. Some of these include co-regulated genes that share the same TAD [27] and clusters of enhancer-promoter interactions observed within TADs [28]. The CCCTC-binding factor (CTCF) has many functions, one of which is to act as an insulator that prevents enhancer-promoter interactions. CTCF also contributes to loop formation and is enriched at certain TAD-boundaries. The disruption of CTCF binding sites present in TAD boundaries through DNA hypermethylation lead to ectopic transcription via interactions across TAD boundaries in cancer [29].

Furthermore, it has been shown that many TAD boundaries are conserved across cell types, with only a small subset of boundaries being cell type specific [24, 30]. TAD boundary regions often coincide with histone modifications associated with regions of gene transcriptional activity, with active enhancer-promoter associated histone modifications such as tri-methylation of Lysines on Histone H3 (H3K4me3 and H3K36me3) and the localisation of essential (housekeeping) genes [23, 24]. CTCF and cohesin were proposed to promote the extrusion of loops, which contributes to TAD formation [31, 32]. According to this model, cohesin extrudes chromatin outwards until it meets barriers in the form of CTCF [31, 32, 33]. Interestingly, the termination of extrusion through CTCF seems to be dependent on the orientation of the protein, as they appear predominantly in convergent orientation [23, 34]. In a Hi-C contact map, these loops anchored at CTCF binding sites correspond to intense, highly localised signals. Although the extrusion model is not yet completely experimentally tested, there is strong evidence supporting it. For example, changes in the orientation of CTCF can change and disrupt chromatin loops and TADs [32, 34]. Furthermore, chromatin loops and TADs are globally lost or reduced upon depletion of cohesin [35, 36, 37].

Chromatin interactions, loops, TADs and A/B compartments can be identified using Hi-C based approaches. Specialised versions of the assay such as promoter capture Hi-C (cHi-C) can detect targeted enhancer-promoter interactions, as well as interaction at the single cell level using single cell Hi-C (scHi-C). These methods need considerable technical expertise, are time consuming and costly. It has also been estimated that to capture chromatin interactions in a cell population at sub-kilobase resolution at least 5-10 billion reads are needed [23]. While recent advances in sequencing technologies make this possible, it is far too expensive to carry out these deep Hi-C experiments. Many models and computational approaches have been developed to predict chromatin interactions and topological domain boundaries. These include an eclectic collection of computational methods including TADLactua [38], PCGS [39], nTDP [40] and HubPredictor [41]. TADLacuta utilised random forest and deep learning methods and integrated histone modifications, CTCF binding and transcription factor motifs predict TAD boundaries with an AUC ROC accuracy 0.867 [38]. In the last five years machine and deep learning technologies have provided revolutionary solutions to many difficult problems in biomedicine [42]. Their utility in genomics [43], biomedical image classification [44], protein structure prediction [45] and many clinical applications [46, 47, 48] have been demonstrated. In this study, we investigate the possibility of using combinatorial histone modification patterns, detected with machine and deep learning approaches to identify both spatial and functional chromatin interactions and architectures.

## Methods

### Cell-lines and data sets

Chromosome conformation capture (Hi-C) and histone modification ChIP-seq data were obtained for three different cell types: IMR90, a human fetal diploid lung fibroblast cell line; K562, a human chronic myelogenous leukemia cell line; and GM12878, a human B-lymphocyte cell line. Hi-C and CTCF ChIP-seq data-sets with the following accessions were used (GSE43070, GSE63525, GSE29611, GSE30263, GSE33213, GSE31477, GSE32465). All data processing steps are visualised by the workflow in Figure S8.

### Histone modification ChIP-seq data

#### ChIP-seq data processing

ChIP-seq data was downloaded from the Roadmap Epigenome project [49] and included fold-enrichment and p-values. A list of available histone modifications, chromatin accessibility and DNA binding proteins for the three cell types in this study can be found in Table S5. CTCF binding profiles for all three cell types were separately obtained from ChIP-seq experiments with accession numbers denoted in Table S1. ChIP-seq reads were mapped to the human reference genome (hg19) using the *BWA aligner (version 0.7.12)* [50]. Low-quality reads (mapping quality *<*20) as well as known adapter contamination were trimmed using *Cutadapt (version 1.10.0)* [51], and reads mapping to the ‘blacklisted’ regions identified by ENCODE [52] were further removed. Average fragment size was determined using the *ChIPQC* Bioconductor package [53], and peak calling was performed with *MACS2 (version 2.1.0)* [54], using fragment size as an extension size (–extsize) parameter.

When computing the signal in a given genomic interval, fold-enrichment values were set to zero where the corresponding p-value was greater than 0.01. The remaining values were binned by taking the mean over a distance of 200 base pairs. We chose this bin size since it is approximately the length of DNA wrapped around a typical nucleosome [55] inclusive of the linker DNA. We then applied an arcsine transformation to the data and divided the resulting data by their mean. The data transformations were carried out to ensure that values were close to 1, since it has been empirically found that neural networks perform better with input values close to 1 [56].

### Hi-C data sets

Hi-C data were obtained in the form of unprocessed fastq files (paired-end Illumina reads) for IMR90 cells from the NCBI Gene Expression Omnibus (GEO)-accession GSE43070[12]. The data set consisted of six biological replicates that were merged for downstream analysis. For K562 and GM12878, the data was obtained from [23], from NCBI GEO, with accession: GSE63525. Due to the size of the files, not all biological replicates were downloaded but only the largest files. Accession numbers are denoted in Table S2.

#### Generation of Hi-C interaction maps

Interaction maps for each data set were generated using *Hi-C-Pro (version 2.11.1)* [57]. The pipeline used the *Bowtie2 (version 2.3.5)* aligner [58] and reads were mapped to the hg19 reference genome, containing chromosomes 1-22 and X. The *in silico* restriction fragment maps were generated for IMR90 using the restriction enzyme *HindIII* whereas maps for the other two cell lines were generated using *MboI*. The interaction matrices were generated for different bin sizes (chromatin loci): 20000, 40000, 80000, 150000, 500000 and 1000000 bps. Different chromatin features are detected at different resolutions, therefore we used bins of 20000 bp resolution for HICRs detection and 40000 bp for TADs.

#### Normalisation of Hi-C interaction maps

The interaction matrices were normalised using the ICE (iterative correction and eigenvector decomposition) method to eliminate systematic biases [59]. The ICE method was applied within the *Hi-C-Pro* pipeline using the Python package *iced (version 0.5.1)*.

#### Significant interactions detection

We used *FitHiC (version 1.3.0, Bioconductor)* [60], a tool for assessing statistical significance of chromosomal contacts with respect to a background, to obtain a list of significant interactions. The original source code of *FitHiC* was altered to consider the interaction matrices from all chromosomes instead of single chromosomal matrices. *FitHiC* returns a p-value and a multiple testing corrected q-value for each interaction. The default parameters were used when running *FitHiC*. Based on number of significant chromatin interactions, we ranked the interacting chromatin loci according to their interaction frequency.

In more detail, we defined significant interactions as those with a q-value below 0.05. Then, for each bin, we summed up the total interaction frequency of significant interactions with other bins. Bins that were in the top 10% were defined as highly interacting chromatin regions (HICRs). Non-HICRs were sampled uniformly from bins with low interaction frequency (lower 70%).

#### TAD boundary detection

*TADbit (version 0.4.28)* [61] was used to call TADs. *TADbit* analyses the contact distribution along the genome and subsequently segments it into TADs. To calculate the position of boundaries between TADs along a chromosome, *TADbit* employs a breakpoint detection algorithm [62]. TADs were called on interaction matrices of 40 kb resolution and only TADs with a *TADbit* score greater than 5 were kept. This ensured that TADs were only called when they spanned more than three bins since smaller regions cannot reliably be called with the breakpoint detection algorithm. TAD boundaries were then defined as either the start or end of adjacent TADs. To create a balanced training set, we randomly chose non-boundaries, equal to the number of TAD boundaries.

#### Identifying biological significance of HICRs and TADs

Functional enrichment analysis of the HICRs was performed using *GREAT (version 3.0.0)* [63]. Overlaps of HICRs and TAD boundaries were calculated and visualized using the R package *ComplexHeatmaps (version 2.3.3)* [64]. TAD boundaries associated with CTCF were identified by selecting those with a CTCF ChIP-seq peak within +/-20 kb of the boundary using *BEDTools (version 2.18)* [65]. The 20 kb window was chosen because this reflects the inherent uncertainty in the exact position of the domain calls due to 40 kb binning. Promoter proximal interactions were mapped to a −5Kb, +1Kb window around TSS using *ChIPpeakAnno (version 3.20.1)* Bioconductor package [66]. The genes were then used in Gene Ontology analysis using the Panther web tool (ver. 15.0) [67].

### Machine Learning Approaches

All learning tasks were designed as binary classification tasks with labels *y* = 1 for positive examples and *y* = 0 for negative examples. The aim was to predict if histone marks in a given region classified as a HICR or not and TAD boundary or non-TAD boundary. We applied the following three supervised machine learning techniques in training to detect HICRs and TAD boundaries and compared them. We used the same training and test set for all three models with a train-test split of 0.8-0.2. All models were implemented in Python *(version 3.7.5)*. Bayesian hyper-parameter optimisation maximising the mean accuracy on the validation set (5-fold cross validation on the training set) was performed using the Python package *hyperopt (version 0.1.2)* [68]. Hyperparameter spaces of all three models can be found in the Supplementary Data, Tables S6 - S8. Predictions were generated on the test set after refitting on the training set.

#### Gradient boosted ensemble of decision trees

The gradient boosting library *XGBoost (version 0.90)* was used for model fitting [69]. The mean over the histone mark signal at the HICRs and TAD boundaries was used as input features. Since the signal for HICRs was very localised, we restricted the region to the extent of the HICR (here 20 kb). For TAD boundaries, we extended the region of the actual TAD boundary (40 kb) on both sides to a total length spanning 200 kb.

#### Logistic regression

The class SGDClassifier from the *scikit-learn library (version 0.21.2)* [70] was used for model fitting. The strength of the regularisation parameter and solver were selected by maximising the accuracy with 5-fold cross-validation. Input features were the same as for XGBoost.

#### Convolutional Neural Network model

A convolutional neural network was constructed in *Keras (version 2.2.4)* with *Tensorflow (version 1.14.0)* as back-end. Here, we did not average the input signal, but used each bin in a window of 200 kb around HICRs and TAD boundaries as one input node to the network. An overview of the architectures can be found in Fig. S9. One convolution block consisted of a convolution layer, batch normalisation before the ReLU activation and a max pooling layer. Before the fully connected layers, we applied dropout. The final classification was done using a sigmoid activation function with a threshold of 0.5. For training, we used the Adam learning algorithm [71] with cross-entropy loss. Most hyperparameters such as the learning rate, number of filters, kernel size, dropout rate were optimized with *hyperopt* (Table S8).

#### Feature importance

The importance of individual features was measured using mean absolute SHAP values (*shap (version 0.32.1)*) of predictions made on the training set.

#### Performance Evaluation

Classification performance of each task is evaluated by the area under the precision-recall curve (PR-AUC) and receiver operating characteristics curve (ROC-AUC). Additionally, we reported accuracy, recall, precision and F1 (harmonic mean of precision and recall) for each task.

#### Regulatory Element Analysis

Genomic annotation information for enhancers (Enhancer-Atlas 2.0 [72], Fantom5 Enhancers [73],GeneHancer [74], SEdb enhancers [75]), super-enhancers (dbSUPER [76], SEdb-super-enhancers), regulatory elements (Ensembl v99 Regulatory Build [77]), genes and TSS (Ensembl v75, Fantom 5), CpG islands, Conservation (UCSC phastCons conservation scores for the human genome hg19 calculated from multiple alignments with other 99 vertebrate species), fragile sites (HumCFS [78]), repeats (microsatellites, repeat-masker elements) and TF clusters (UCSC table browser) [79] were downloaded from respective databases. Enrichment analysis was performed with GAT (Genomic Association Test), testing the number of overlapping segments with 3000 permutations [80].

## Results

### Overview

Three different cell lines were used in our analyses; these included GM12878 transformed B lymphocytes, K567 myelogenous leukaemia cells and IMR90 diploid fetal fibroblasts. Fig. 1 gives an overview of the methodology used. Features were extracted from ChIP-seq derived histone modification data. Details of the data sets, data cleaning, processing and feature extraction is described in Methods. Fold-enrichment values and p-values for all available histone markers and CCCTC-binding factor (CTCF) binding profiles were obtained for all three cell types, see Table S5 for a detailed list. Signal regions binned by aggregating over a distance of 200 base pairs were used to extract features (Fig. 1a). We chose this bin size since it is approximately the length of DNA wrapped around a nucleosome [55] plus its linker DNA.

**Fig. 1:**
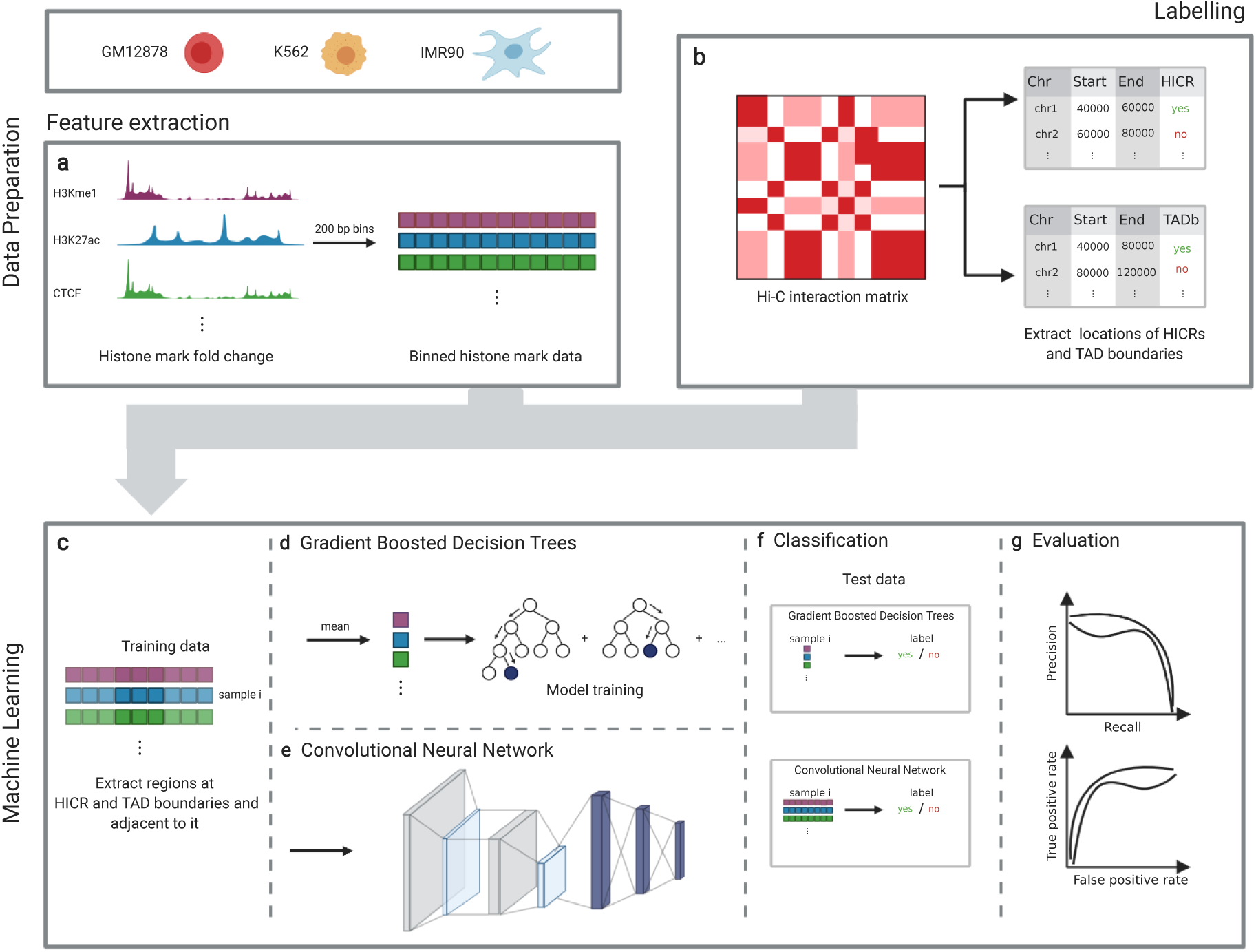
Overview of workflow. All analysis steps were performed for three different cell types – IMR90, GM12878 and K562. **a-b**, Data Preparation. **a**, For feature creation, fold-change values for all available histone marks were aggregated over 200 bp bins, the typical length of DNA spooled over nucleosomes with linker DNA. **b** As targets, highly interacting chromatin regions (HICRs) and TAD boundaries were extracted from normalised Hi-C interaction matrices. For HICRs, a resolution of 20 kb was used, for TAD boundaries 40 kb. All interacting loci were then labelled as either HICR or TAD boundary and non-HICR or non-TAD boundary to construct binary classification tasks. For training, we sampled a subset of interacting loci with negative label, to create a balanced set. **c-f**, Machine Learning. **c** For each target (HICR or TAD boundary), bins in a window adjacent to it were extracted. 80% of the data was used for training, with 5-fold cross validation. The remaining 20% were used for the final evaluation. **d,e** A binary classifier to predict HICRs and TAD boundaries was trained on the extracted labels and features. Both gradient-boosted decision trees and convolutional neural networks were evaluated. In the first case, the histone mark signal was averaged to have one feature per histone mark. **f**, Trained models were used to classify the left-out test data. **g**, Evaluation. The models were evaluated on the basis of precision-recall curves and receiver operating characteristics (ROC) curves.

Similarly, chromatin interactions were extracted from iced-normalised interaction matrices from Hi-C data. TAD boundaries were called using these matrices and statistically significant boundaries were identified. Highly Interacting Chromatin Regions (HICR) were identified. A balanced training set was defined for TAD boundaries and HICRs targets from normalized Hi-C contact maps for three different cell types – IMR90, GM12878 and K562, Figure 1b. Machine and deep learning approaches were used to create a binary classification model, which classifies a region in the genome as HICR or TAD boundary (labels) from the underlying histone mark signal (features). The process of label and feature creation and model training is illustrated in Fig. 1c–g.

### Properties of HICR, TAD boundary targets and histone mark features

#### Defining highly interacting chromatin regions (HICR)

HICRs were defined by significant interactions from the iced-normalized interaction count matrix, identified by *FitHiC* [60], Fig. 1b. For each chromatin locus, we counted the number of statistically significant interactions, which showed a similar long tailed distribution in all three cell types (Fig. 2a for GM12878 and Fig. S1a–b for IMR90 and K562 respectively). Across all three cell types, 30% of all loci showed a high significant interaction frequency, with the remaining 70% showing very few interactions. Interestingly, although the distributions were similar, the frequency differed between cell types. In K562 cells, the top 10% of interacting regions showed more than 3700 interactions per locus, while IMR90 cells showed a minimum of 890 interactions, and GM12878 cells showed an intermediate value of around 1700 interactions. The differences here could be both technical (the IMR90 Hi-C data-set has lower resolution) or biological in origin.

**Fig. 2:**
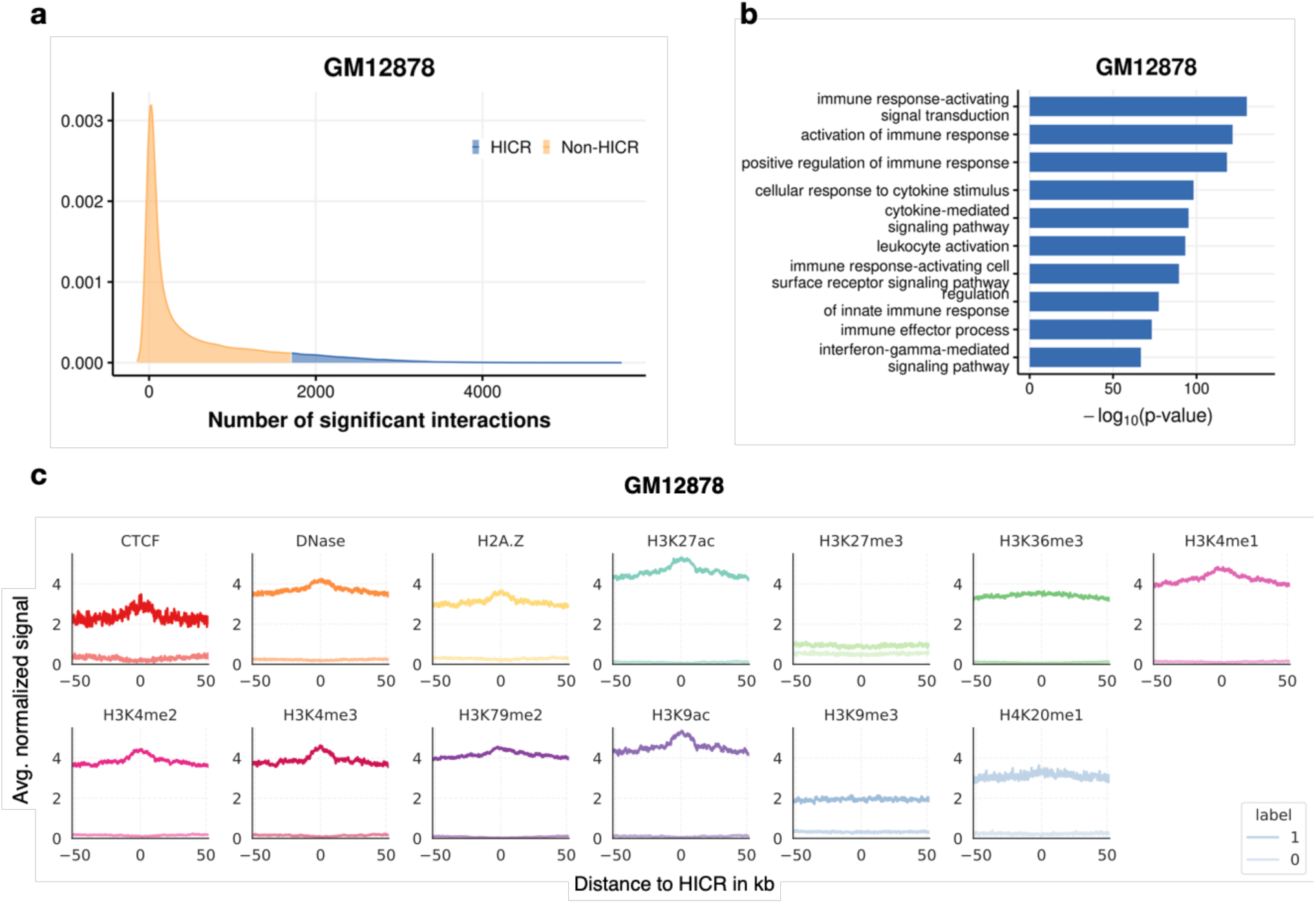
Inspection of HICRs. **a**, The distribution of significant counts for the GM12878 cell line. The top 10% of interactions are shaded in blue, which were defined as Highly Interacting Chromatin Regions (HICRs). **b**, Functional enrichment analysis for HICRs in the GM12878 cell line. The top 10 terms are displayed, ordered by the negative log10, which is displayed on the x-axis. In GM12878, a B-lymphocytes cell line, terms related to immune response were highly enriched. **c**, Spatial pattern of the histone marks found in a window of 100 kb adjacent to the HICRs. The signal was averaged over all samples. The average signal over non-HICRs is illustrated in a lighter colour. In general, histone marks that are linked to transcriptional activity (acetylations, H3K36me3, H3K4me1) had an increased signal.

#### Over-representation analysis of HICRs

To determine if loci with a high interaction frequency either have a role in maintaining chromatin spatial structure or a functional role in transcriptional regulation, or both, we investigated the relevance of these highly interacting loci. The enrichment of these regions for biologically significant genomic elements or annotation were explored. We divided the set into two; the top 10% and next 20%. Since they formed the set of most highly interacting genomic regions of each cell type, we scrutinised these regions for their contribution to cell-specific gene regulation.

Functional enrichment analysis using GREAT and Panther web-tools provided insights into the biological functions of the highly interacting chromatin regions. Indeed, genes nearby the interacting loci in the top set were significantly enriched for cell type-specific processes (Fig. 2b and Fig. S1c–d). Interacting chromatin loci of the IMR90 cell line, a human fetal lung cell line, were enriched for developmental processes related to the mesodermal origin of lung fibroblasts, such as mesoderm development (P-value = 5.5× 10^−46^) or positive regulation of angiogenesis (P-value = 4.7× 10^−54^). In GM12878, a B-lymphocyte cell line, terms related to immune response were highly enriched, e.g.positive regulation of immune response (P-value = 2.4 × 10^−121^). Leukocyte mediated immunity, as well as response to endoplasmic reticulum stress, were enriched in K562 interacting loci, from a human chronic myelogenous leukaemia cell line.

Endoplasmic reticulum (ER) is involved in protein folding and assembly, Ca^2+^ homeostasis, and lipid biosynthesis. Multiple stress conditions can trigger ER stress leading to an unfolded protein response and is commonly observed in both acute and chronic myelogenous leukaemia [81]. In the next 20% group (10% – 30%) we found more general cellular processes such as regulation of metabolism, signalling and protein modifications, regulation of transcription, cytoskeleton and organelle organization. These could be interactions involved in regulating essential cellular processes common between cell types. As a control, we also analysed chromatin loci at the lower end of the distribution, with no significant results for functional enrichment. Since we were interested in investigating cell type specific properties, we defined the top 10% as our highly interacting chromatin regions (HICR). For the set of non-HICRs, we uniformly sampled an equal number from the low-interacting loci (bottom 70%).

#### Histone modification enrichment in HICR flanking regions

To explore if particular histone modification signatures were correlated with HICRs, we examined the patterns of histone modifications at each HICR and its neighbouring regions in a 100 kb window. A distinct signal is evident in the histone modifications at the HICRs (Fig. 2c). For comparison, we illustrate the background signal found in regions that were not identified as HICRs in a lighter colour. We found high enrichment for CTCF (suggesting local loop formation) and activating histone modifications (H3K27ac, H3K4me1, H3K4me3) associated with active enhancer and promoter regions. In addition, regions containing transcriptionally active gene bodies (H3K36me3) and DNA damage response associated modifications (H2AK5Ac, H2BK120Ac, H4K20me1) were enriched compared to non-HICR control regions. DHS assays in GM12878 showed that these active regions were also nucleosome free open regions. Similarly, heterochromatin modifications such as inactivating H3K9me3 and H3K27me3 marks showed very little enrichment compared to non-HICR regions (visualised in a lighter colour), however these heterochromatin effects were variable across the cell types with slight enrichment of H3K27me3 in K562, associated with facultative repression of promoter rich regions (Fig. 2c and Fig. S2a and b).

#### TAD sizes are very similar between the three cell types, are enriched for CTCF at boundaries and show little enrichment of most histone modifications

We checked the distribution of sizes of the TADs to verify if they were on a reasonable length scale (Fig. 3a). The interquartile range (IQR) of TADs was between 0.25 and 0.75 Mb for all three cell types with a median size of 500 kb, which is in agreement with [23]. A few very large TADs were predicted that could most likely are miscalled boundaries by *TADbit*. Fig. 3c and Fig. S3a and b show the spatial pattern of histone marks in a 300 kb window around the TAD boundaries. It is striking that compared to that observed in HICRs the signal was a lot weaker and the peaks broader, however, there were some histone modifications that were enriched at TAD boundaries. Most importantly, CTCF, a marker of TAD boundaries, was enriched. Additionally, markers associated with active promoters and genes showed a stronger signal (H3K36me3 in all three cell types and H3K4me3 in IMR90 cells) which is distinctly different to repressive marks that were not enriched or even depleted in boundary regions (H3K9me3 and H3K27me3). Again, this is in agreement with findings that TAD boundaries correlate with histone modifications that are associated with active gene regions and are often marked by the CTCF-binding sites [23].

**Fig. 3:**
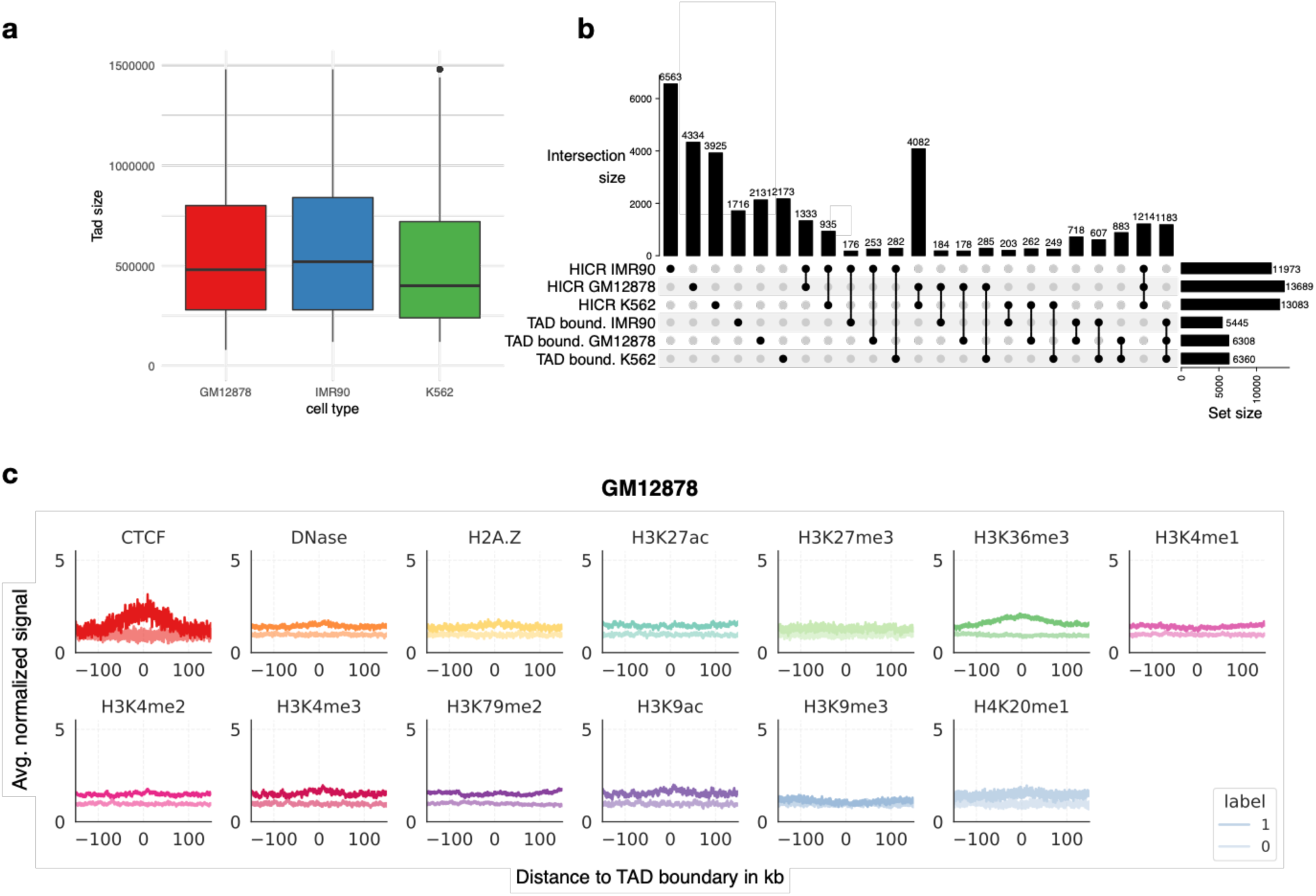
Inspection of TAD boundaries. **a**, Distribution of the sizes of identified TADs for all three cell types. The interquartile range of TAD sizes was between 0.2 Mb and 0.75 Mb with a mean of 0.5 Mb.**b**, UpSet plot showing the intersection of identified HICRs and TAD boundaries. Intersections of more than two sets are not shown except for the intersection of all three HICR sets and TAD boundary sets. IMR90 had a high percentage of unique HICRs (≈ 55%). The 30% overlap between HICRs in K562 and GM12878, is most likely attributed to the fact that both cell types are derived from immune cells. Roughly 10% of all HICRs were common to all three cell types. 20% of all TAD boundaries were shared across all three cell types, whereas only 30% could be attributed to a single cell type. The number of inter-HICR and TAD boundary overlap was very small. **c**, Spatial pattern of the histone marks found in a window of 300 kb adjacent to the TAD boundaries. The signal was averaged over all samples. The average signal over non-TAD boundaries is illustrated in a lighter colour. Compared to the HICRs, the signal was a lot weaker and the peaks broader. Most importantly, CTCF and H3K36me3 – both well-known markers of TAD boundaries – were enriched.

#### Overlap between HICRs and TAD boundaries

Fig. 3b compares HICRs and TAD boundaries between cell types. IMR90 showed a high percentage of unique HICRs (≈ 55%). In contrast, only a third of all HICRs were unique for GM12878 and K562. The 30% overlap between HICRs in K562 and GM12878, is most likely attributed to the fact that both cell types are derived from immune cells and have similar highly interacting regions contributing to their cell type specificity. Roughly 10% of all HICRs were common to all three cell types. Looking at TAD boundaries, there was no such difference between cell types. 20% of all TAD boundaries were shared across all three cell types, whereas only 30% could be attributed to a single cell type. The number of inter-HICR and TAD boundary overlap was very small.

### Binary classification of HICRs and TAD boundaries

We developed and evaluated three different model types – logistic regression, gradient boosted decision trees (XGBoost) and a convolutional neural network (CNN). Here we present the results from the XGBoost and the CNN. Logistic regression performed surprisingly well, but overall did not outperform the XGBoost or CNN model (see Table S9 and S10).

#### XGBoost model

We trained two binary XGBoost models per cell type, one for HICRs and one for TAD boundaries. Each model learnt to classify HICR/TAD boundary vs non-HICR/non-TAD boundary using combinatorial histone mark signatures. As input, we used the mean of the histone mark signal over a certain region at the HICRs and TAD boundaries, see Fig. 1c–d. We evaluated different window sizes, ranging from 20 kb to 300 kb. For HICRs we found that a very small window of 20 kb produced the best results (which is equivalent to the Hi-C data resolution), TAD boundaries required a broader region of 200 kb. Additionally, we trained the models with all histone marks that were available for each cell type and also with only the consensus set of histone modifications available for all three cell types (Table S5) for better comparability of the different combinatorial effects of histone modifications.

The performance of the boosted tree classifier for GM12878 is shown in Fig. 4 using both ROC curves and precision-recall (PR) curves and in Fig. S4 for the other two cell lines. Additional metrics are shown in Table 1 and 2. Overall, there is very good performance across all cell types when predicting HICRs, with only a small range between the best performing model (ROC-AUC 0.999 and PR-AUC 0.999) and the worst performing model (ROC-AUC 0.981 and PR-AUC 0.977).

**Table 1:**
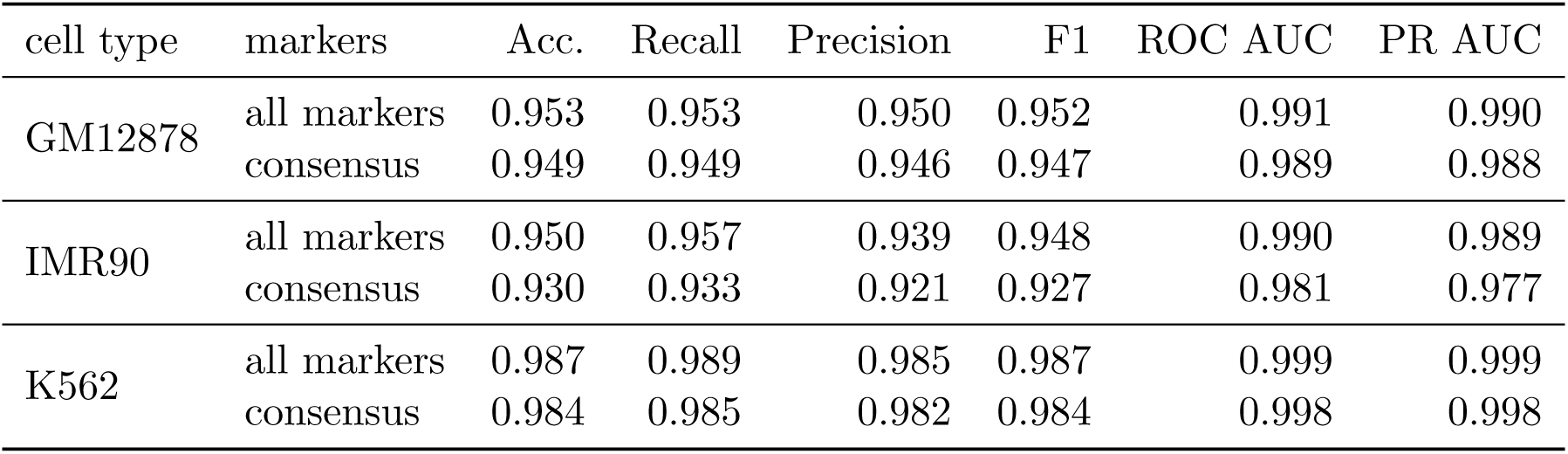
HICR: Metrics used for evaluation of XGBoost models for each cell type.

**Table 2:** TAD boundaries: Metrics used for evaluation of XGBoost model for cell type.

| cell type | markers | Acc. | Recall | Precision | F1 | ROC AUC | PR AUC |
| --- | --- | --- | --- | --- | --- | --- | --- |
| GM12878 | all markers | 0.715 | 0.818 | 0.670 | 0.737 | 0.774 | 0.722 |
|  | consensus | 0.706 | 0.807 | 0.664 | 0.728 | 0.772 | 0.724 |
| IMR90 | all markers | 0.704 | 0.808 | 0.663 | 0.728 | 0.770 | 0.713 |
|  | consensus | 0.697 | 0.790 | 0.659 | 0.719 | 0.762 | 0.708 |
| K562 | all markers | 0.707 | 0.795 | 0.683 | 0.735 | 0.757 | 0.715 |
|  | consensus | 0.702 | 0.796 | 0.678 | 0.732 | 0.757 | 0.718 |

**Fig. 4:**
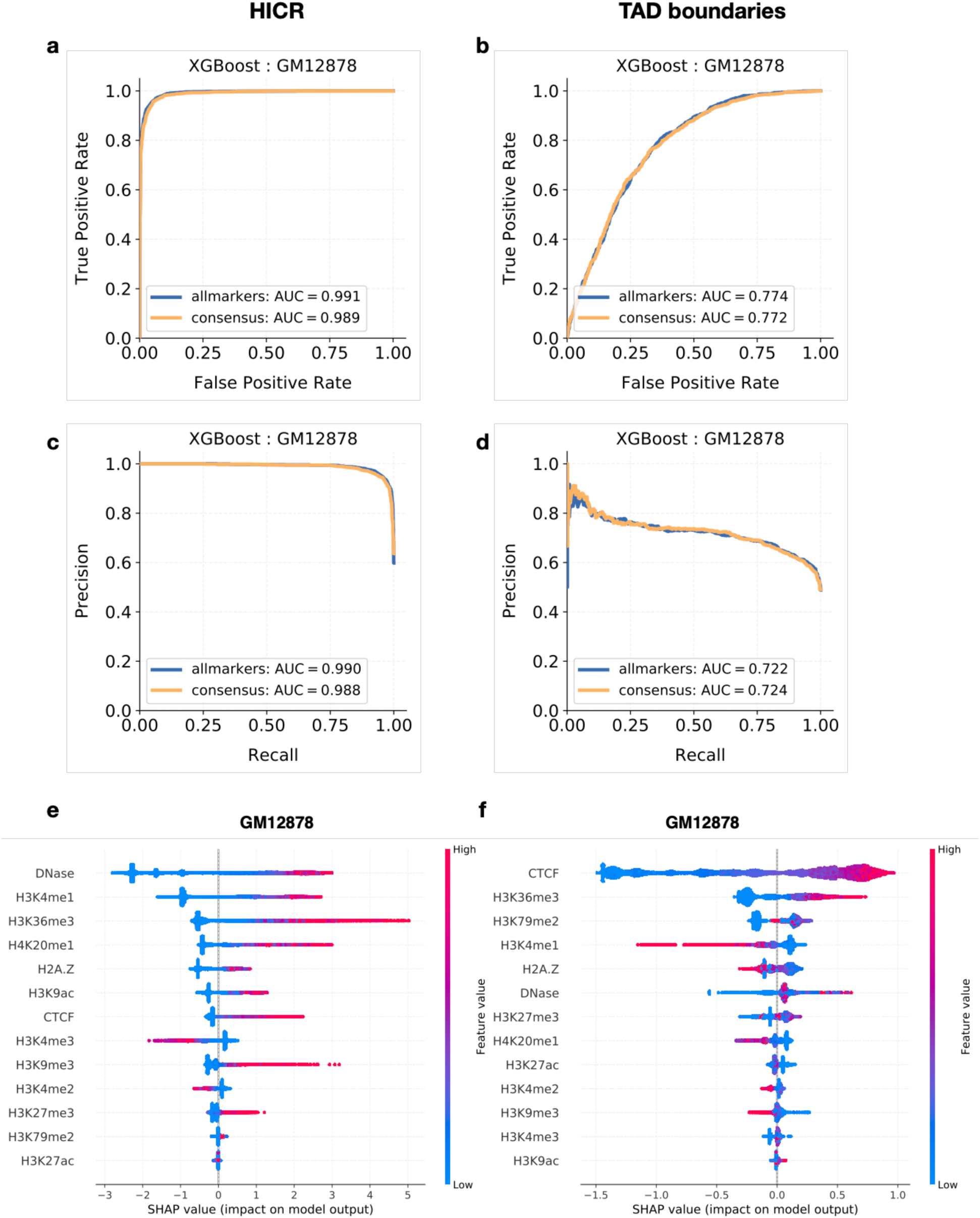
XGBoost performance and Feature importance on GM12878. **a-b** Receiver-operating characteristic curve and **c-d** precision-recall curve for the binary classification task of predicting HICRs (**a, c**) and TAD boundaries (**b, d**) from histone mark data. Models were trained once using all available histone marks for each cell type (blue) and using only the consensus set (orange). **e-f** Histone marks ordered by mean absolute SHAP values in HICRs (**e**) and TAD boundaries (**f**). On the y axis, the violin plot shows the full distribution of the SHAP values for each feature. The dot plot in the foreground shows a color coding of the actual value of the feature, resulting in the SHAP value as indicated on the x axis.

Classification of TAD boundaries proved to be more difficult with the best performing model getting a test performance of 0.774 (ROC-AUC) and 0.722 (PR-AUC). One explanation here could be the lack of significant histone mark signals at and adjacent to the TAD boundaries (Fig. 3c).

Interestingly, the use of all histone marks compared to the consensus set did not have a large effect on model performance. The biggest increase in test set performance could be seen for IMR90, which had most histone marks available (27 in total), which indicates that model performance might increase when taking additional histone marks into account that were not available for the two other cell types.

#### Inspection of model features

We used SHAP values to get an insight into which histone marks are predictive for both HICRs and TAD boundaries. In Fig. 4e, we list the features of the HICR model for GM12878 sorted by mean absolute SHAP value and show the distribution across all predictions. Additionally, sample points are coloured according to their feature values. An increased signal of well-known markers for active regulatory elements such as DNase Hyper-sensitivity Assay (DHS), H3K4me1 and H3K36me3 are strong indicators for the presence of a HICR. On the contrary, higher values of H3K4me3 give lower probability to the positive class. The other two cell types show similar distributions (Fig. S5) with a few striking differences. In IMR90 (Fig. S5a), the acetylation of H2AK5 has the highest feature importance, a mark that is linked to active regions [82]. This could also explain the observed increase in performance when using the full model (all markers) compared to only the consensus set. Also, in both IMR90 and K562, more importance is given to the repressive mark H3K27me3, indicating that we might be capturing both active and inactive (facultative heterochromatin) regulatory elements.

Fig. 4f and Fig. S5b and d show the distribution of SHAP values for the prediction of TAD boundaries. TAD boundaries are often marked by CTCF [24] and its presence has the biggest influence on model prediction in all three cell types. In GM12878, both H3K36me3 and H3K79me2, that have previously been found to be associated with TAD boundaries [23], also have a high feature importance. Interestingly, H3K4me1 has an anti-correlated relationship with model prediction, indicating that TAD boundaries are depleted for this enhancer associated mark. In IMR90 and K562, H3K9me3 ranks among the top four most important features with an anti-correlated relationship. In both cases the feature analysis has shown that the models pick up relationships between the chromatin feature and histone marks that were not apparent by merely looking at the average signal.

#### Convolutional Neural Network

With the aim of creating a more robust model, especially for TAD boundary predictions, we designed two convolutional neural networks, with the idea that it would better capture patterns within the histone mark signal (see Methods and Supplementary Data for the detailed architecture and hyperparameter space). The network was provided with a sequence of binned histone mark data in a window of 200 kb adjacent to HICRs and TAD boundaries. Fig. 5a–d, Fig. S6 and Tables 3 and 4 show the performance on the test set. Performance of the CNN at predicting TAD boundaries was improved from a mean ROC-AUC of 0.765 to 0.784 and average PR-AUC of 0.717 to 0.755. For HICRs performance was comparable to the XGBoost model. The mean ROC-AUC over all six models was slightly improved from 0.991 to 0.997 and mean PR-AUC from 0.990 to 0.997.

**Table 3:** HICR: Metrics used for for evaluation of CNN models for each cell type.

| cell type | markers | Acc. | Recall | Precision | F1 | ROC AUC | PR AUC |
| --- | --- | --- | --- | --- | --- | --- | --- |
| GM12878 | all markers | 0.989 | 0.992 | 0.984 | 0.988 | 0.999 | 0.999 |
|  | consensus | 0.981 | 0.995 | 0.968 | 0.981 | 0.998 | 0.998 |
| IMR90 | all markers | 0.978 | 0.975 | 0.980 | 0.977 | 0.997 | 0.997 |
|  | consensus | 0.958 | 0.971 | 0.942 | 0.956 | 0.988 | 0.987 |
| K562 | all markers | 0.995 | 0.993 | 0.996 | 0.995 | 0.999 | 0.999 |
|  | consensus | 0.996 | 0.996 | 0.996 | 0.996 | 1.000 | 1.000 |

**Table 4:** TAD boundaries: Metrics used for evaluation of CNN models for each cell type.

| cell type | markers | Acc. | Recall | Precision | F1 | ROC AUC | PR AUC |
| --- | --- | --- | --- | --- | --- | --- | --- |
| GM12878 | all markers | 0.733 | 0.797 | 0.709 | 0.750 | 0.803 | 0.784 |
|  | consensus | 0.723 | 0.836 | 0.684 | 0.752 | 0.787 | 0.765 |
| IMR90 | all markers | 0.733 | 0.810 | 0.706 | 0.754 | 0.806 | 0.776 |
|  | consensus | 0.725 | 0.841 | 0.687 | 0.756 | 0.794 | 0.759 |
| K562 | all markers | 0.697 | 0.710 | 0.693 | 0.701 | 0.757 | 0.722 |
|  | consensus | 0.694 | 0.838 | 0.651 | 0.733 | 0.758 | 0.724 |

**Fig. 5:**
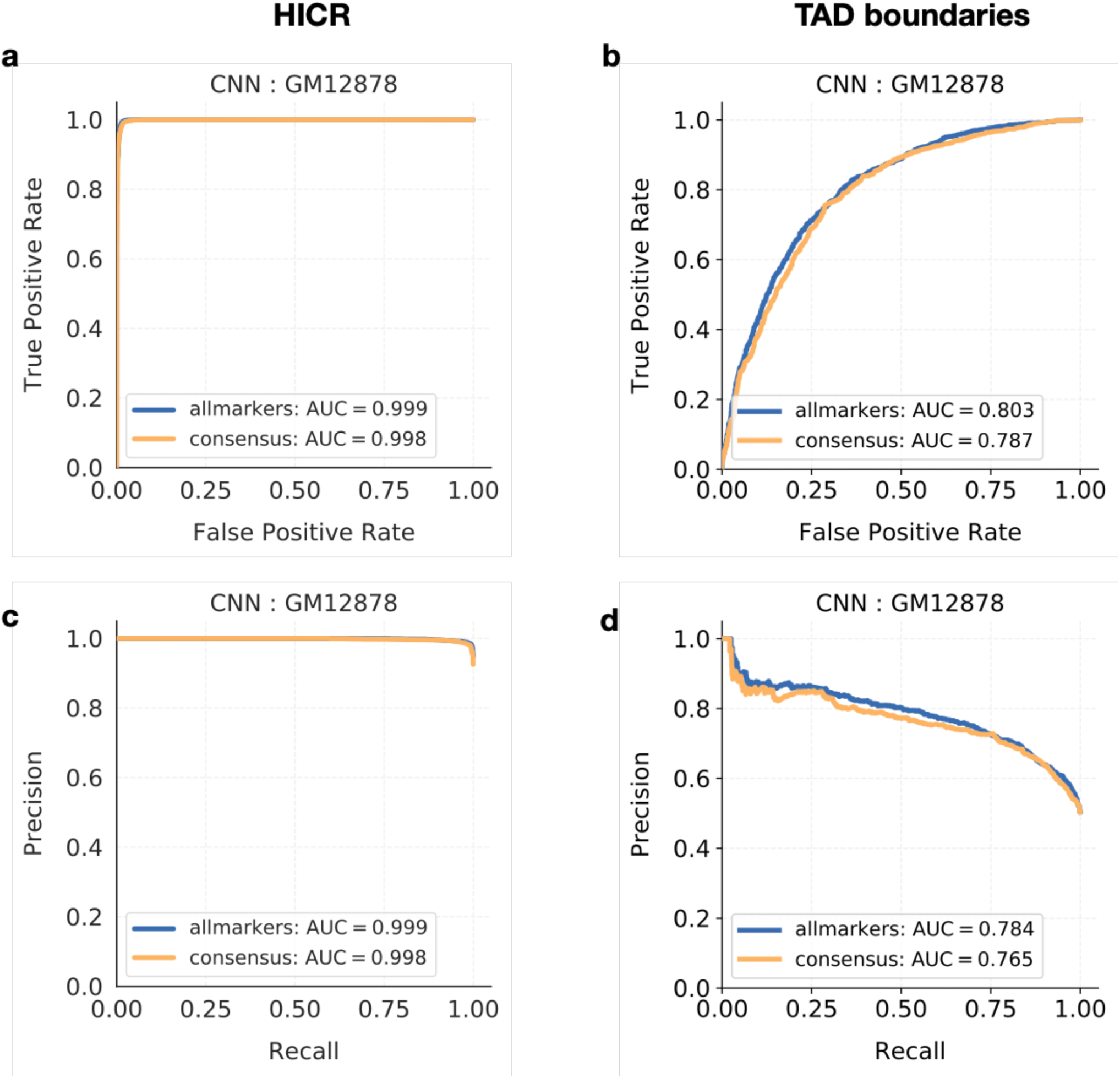
CNN performance on GM12878. **a-b** Receiver-operating characteristic curve and **c-d** precision-recall curve for the binary classification task of predicting HICRs (**a, c**) and TAD boundaries (**b, d**) from histone mark data. Models were trained once using all available histone marks for each cell type (blue) and using only the consensus set (orange). CNNs outperformed the XGBoost model.

#### Transfer models to assess model generalisability across cell types

We found that CNNs trained on cell type specific data perform better than conventional machine learning models at predicting HICRs and TAD boundaries from histone mark data. Building on that, we examined if that was a phenomenon that could be attributed to the cell type specific training data given to the network or if histone marks adjacent to our targets experienced cell type independent patterns. For this, we tested the performance of a model trained on one cell type and tested on data from a different cell type. Since we did not have all histone marks for every cell type, we compared performance using the consensus set. Fig. 6a–d, Fig. S7 and Tables 5–6 show the results.

**Table 5:** HICR: Different metrics for evaluation of CNN model tested on all three cell types.

| cell type trained | cell type tested | Acc. | Recall | Precision | F1 | ROC AUC | PR AUC |
| --- | --- | --- | --- | --- | --- | --- | --- |
| IMR90 | IMR90 | 0.958 | 0.971 | 0.942 | 0.956 | 0.988 | 0.987 |
|  | GM12878 | 0.954 | 0.928 | 0.976 | 0.951 | 0.989 | 0.989 |
|  | K562 | 0.952 | 0.920 | 0.984 | 0.951 | 0.994 | 0.995 |
| GM12878 | IMR90 | 0.928 | 0.887 | 0.959 | 0.922 | 0.983 | 0.982 |
|  | GM12878 | 0.981 | 0.995 | 0.968 | 0.981 | 0.998 | 0.998 |
|  | K562 | 0.969 | 0.946 | 0.991 | 0.968 | 0.993 | 0.994 |
| K562 | IMR90 | 0.906 | 0.831 | 0.969 | 0.895 | 0.983 | 0.982 |
|  | GM12878 | 0.948 | 0.989 | 0.911 | 0.948 | 0.991 | 0.989 |
|  | K562 | 0.996 | 0.996 | 0.996 | 0.996 | 1.000 | 1.000 |

**Table 6:**
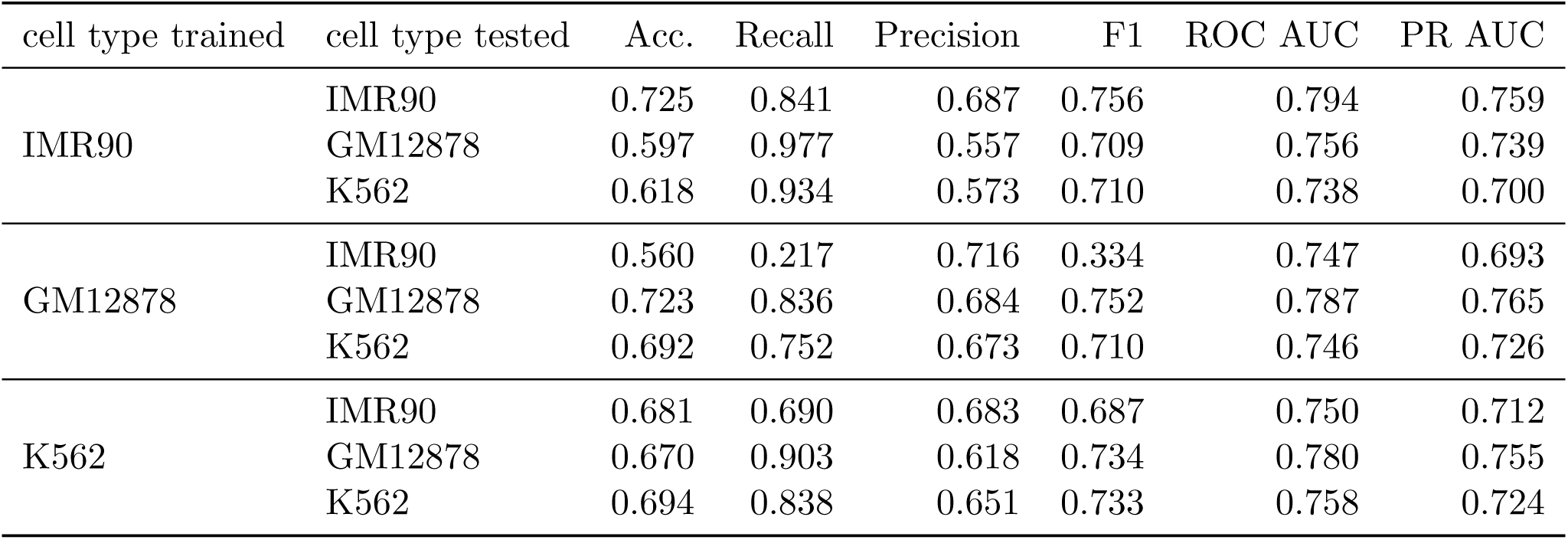
TAD boundaries: Different metrics for evaluation of CNN model tested on all three cell types.

**Fig. 6:**
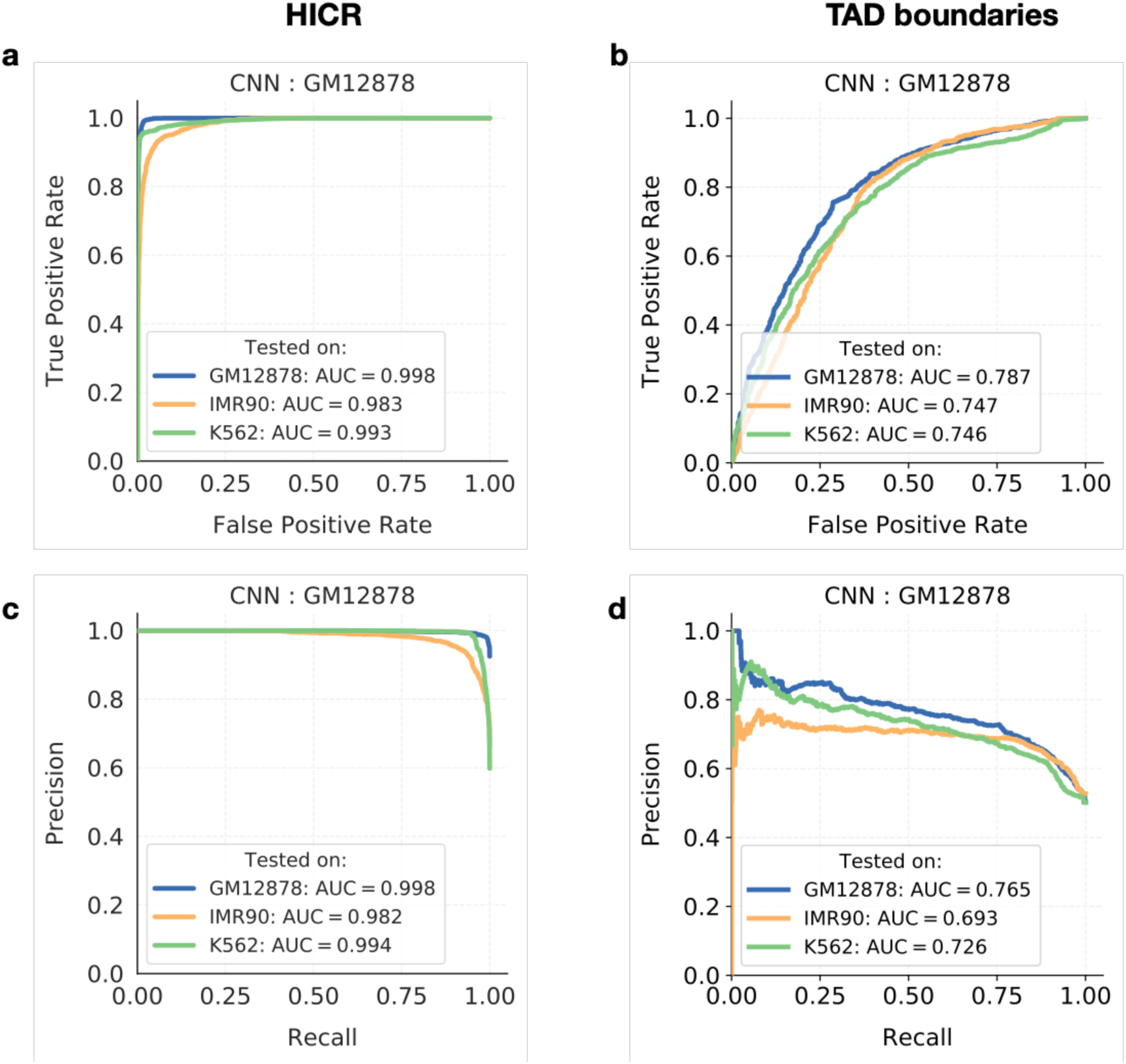
Cross-cell performance on GM12878. **a-b** Receiver-operating characteristic curve and **c-d** precision-recall curve for the binary classification task of predicting HICRs (**a, c**) and TAD boundaries (**b, d**) in all three cell types from models solely trained on GM12878 data. Models were trained using only the consensus set.

Interestingly, the performances are surprisingly good with all AUC values close to the evaluation on the same cell type. Most models perform 1-2% worse on predicting HICRs and TAD boundaries in other cell types. There is one exception, in the case of predicting TAD boundaries in IMR90 from the model trained on GM12878, where the performance significantly decreases. Even though the ROC-AUC and PR-AUC are comparable to testing on the same cell type (ROC-AUC: 0.787 vs. 0.747 and PR-AUC: 0.765 vs. 0.693), we get low accuracy and F1 score (0.56 and 0.334, respectively). This is mainly due to the low recall (0.217), which means that many TAD boundaries are misclassified as non-TAD boundaries. Conversely, when predicting TAD boundaries in GM12878 from models trained on IMR90, accuracy is also not high (0.597). Overall, however, these findings suggest that even though we found that HICRs and TAD boundaries vary across cell types, there might be a more general and cell type invariant pattern of histone modifications, marking these chromatin features.

## Discussion

Precise regulatory mechanisms are essential in controlling transcription and translation of multicellular organisms, where dysregulation of gene expression can potentiate pathological disease states. Our understanding of how chromatin interactions and spatial genome organisation contribute to transcriptional regulation and DNA replication has vastly improved due to the developments in chromosomal conformation capture methods such as 4C, 5C and various Hi-C technologies. Notably, Hi-C interaction maps provide a means to understand how three-dimensional genome organisation and chromatin interactions can affect gene regulation in both normal and disease states [83]. However, these methods are technically challenging, time consuming, have many associated biases and are costly. Altogether, it is a motivating driver for developing computational methods that can predict functional interactions and chromatin architectures as those detected by Hi-C using more easily produced epigenetic data. Patterns of epigenomic activity obtained from ChIP-seq experiments that detect post-translational histone modifications are one such data type. In this study, we demonstrate the utility of histone modification data together with machine and deep learning approaches to develop models that predict chromatin interactions and domain boundaries.

A number of models and computational approaches have previously been produced to predict or identify domain boundaries and interacting regions. Henderson et al., used deep learning methods to predict TAD boundaries using high resolution Hi-C in *Drosophila melanogaster* [84], highlighting the use of deep neural networks applied to chromatin biology. Likewise, Hong et al., [39] developed a population greedy search algorithm (PGSA) to identify boundary enriched motifs and concluded that in addition to CTCF, other TFs such as ZNF143, ZNF274, YY1, SP1 and chromatin accessibility (DHS) as well as epigenetic modifications (H3K36me3), gene associated features (TSS) and transcriptional complexes (RNA PolII) all improved identification of TAD boundaries. Similarly, Sefer and Kingsford developed a convex semi-nonparametric approach called nTDP, based on Bernstein polynomials to explore the joint effects of histone markers on TAD formation [40]. Furthermore, Huang et al., [41] used a statistical approach known as Bayesian Additive Regression Trees (BART) to develop HubPredictor, which detects interacting regions or “hubs” and TAD boundaries. They used a 300 kb window around signal regions and averaged reads for each histone modification. Their prediction ROC AUC was 0.869 for interactions and 0.774 for TAD boundaries. While their models built with averaged cell type data performed well for TAD prediction, cell type specific data was needed for interaction prediction. In their model, CTCF was the most predictive feature for domain boundaries, while H3K4me1 and H3K27me3 were important in interaction prediction. Huang et al., [41] speculate that these combinatorial patterns may be involved in mediating chromatin interactions. However, it is also possible that histone modifications merely demarcate regulatory elements that are involved in functional interactions or spatial genome organisation and are selected by chromatin interacting proteins such as remodelers and transcription factors in forming interactions.

The studies previously described used various data types (including histone modifications) to detect or predict either chromatin interactions or topological features. The approaches used had varying rates of success, and most were restricted to specific species or cell types. We hypothesised that machine and deep learning methods would be more efficient at producing cell type independent models with increased predictive performance. Using Hi-C data from three cell types – IMR90, GM12878 and K562 – we constructed normalised interaction count matrices from which we identified the locations of highly interacting chromatin regions (HICRs) and topologically associated domain (TAD) boundaries as targets of a classification task. We determined that the top 30% of all interactions from Hi-C data are high frequency interactions. Gene ontology enrichment analysis of HICR regions in the three cell types demonstrated that the top 10% of interaction regions contain genes involved in cell type specific functions and the next 20% of interacting regions are enriched for genes contributing to general cellular functions. We tested three different machine learning approaches, out of which gradient boosted decision trees (XGboost) and deep convolutional neural networks (CNN) successfully predicted two important chromatin architectures – HICRs and TAD boundaries – from epigenomic data. The models were subsequently trained on a well studied collection of histone modifications and CTCF ChIP-seq data. Both machine and deep learning approaches demonstrated good test set performance, with the CNN yielding better results in the final evaluation. Whilst we obtained almost perfect test set performance (ROC-AUC 0.999 and PR-AUC of 0.999) when predicting HICRs, TAD boundaries proved more difficult to classify (ROC-AUC of 0.803 and PR-AUC of 0.784). The use of all features compared to just a consensus subset of histone modifications available for all three cell types (Table S3) made little difference. Importantly, models trained on one cell type could predict HICRs and TAD boundaries using only histone modifications from the other two cell types with comparable high-performance. This clearly indicates that certain specific histone modification signatures are associated with both HICRs and TAD boundaries.

We hypothesized that the highly interacting chromatin regions may contain regions contributing to tissue specific regulatory elements or chromatin architecture. In all three cell types, enrichment of both active enhancer (H3K27Ac, H3K4me1), promoter (H3K4me3, H3K4me2), and gene body transcriptional activity (H3K36me3) associated histone modifications suggests that HICRs are enriched for active enhancer-promoter regions as well as transcriptionally active genes. Indeed, we found HICRs are highly enriched for known enhancers, super-enhancers, CpG islands, gene transcription start sites and TF bound regions (Table S3 and S4). Further, we detected enrichment of loop–forming CTCF in HICR regions, increased chromatin accessibility (detected with DNase hypersensitivity assay), enrichment of histone modifications such as H4K20me1 which is associated with transcriptional activation (in GM12878 and K562), and enriched H2A.Z histone proteins (in GM12878 and K562). The latter is a histone that when included into nucleosomes that are proximal to the TSS, correlates with higher gene expression [82, 85]. Conversely, depletion of repressive heterochromatin associated modifications (H3K9me3, H3K27me3) is observed, and there is cell type specificity in these changes. Interestingly, moderately increased activity of the polycomb repressive facultative heterochromatin modification H3K27me3 is seen in HICR regions of the K562 cancer cell line, and in GM12878 cells moderately high levels of constitutive heterochromatin modification H3K9me3 is detected. Our analysis indicates that HICRs may be composed of a heterogeneous collection of active, inactive, poised and primed enhancer-promoter regions. Thus, multiple lines of evidence corroborate that HICRs are enriched for enhancer-promoter regions and actively transcribed genes.

Comparison of genomic locations of all HICRs, showed cell type specificity of the highly interacting regions. The fibroblast cells (IMR90) had less than 20% common HICR regions with the other two cell lines, while K562 and GM12878 (both with an immune origin) contained 40% overlap between HICR regions. In contrast, TAD boundaries were more common with 30–50% shared between cell types. There was very little overlap between HICRs and TAD boundaries between cell types suggesting that DNA regions involved in transcriptional regulation and genome topology organisation are mostly distinct, even though detectable using the same collection of epigenetic modifications. The enrichment of active enhancer–promoter interactions in these HICR regions may contribute to coordinated gene expression programs. These regions are reminiscent of coordinately expressed genes that co-localise in the nucleus forming “transcriptional factories” [86]. The observation of polycomb mediated facultative heterochromatin associated modifications in HICRs is suggestive of specific silencing of regulatory regions and associated genes. These facultative heterochromatin clusters are also known to bring together distal chromatin regions [87]. Formation of DNA-RNA hybrid structures known as R-loops on TSS proximal regions of transcribed genes [88] and chromatin loop anchors lead to fragile sites and double stranded DNA breaks [89]. We observed an increase of fragile regions and enrichment of DNA repair associated histone modifications, similar to previously reported aggregates of repair assemblies associated with double stranded breaks [90]. Thus HICRs may consist of enhancer-promoter interactions contributing to transcriptional factories, targeted silenced regions and regions of repair assemblies.

Interpretability and explainability of machine learning models are a desired characteristic as they provide model transparency. We utilised the SHAP (SHapley Additive exPlanations) method developed by Lundberg and Lee to assess contribution of individual features [91]. SHAP contributes to the interpretability of models by using Shapley values from game theory. The SHAP analysis revealed that the enhancer modification H3K4me1, transcriptional activation modification H4K20me1, and gene body and exon defining modification H3K36me3, which may also be involved in splicing as well as preventing runaway transcription [92, 93, 94], contribute the most to HICR prediction. The SHAP analysis identified CTCF and H3K36me3 to have high predictive power for TAD boundaries, which are well-known associations [23]. However, some importance was also given by SHAP to the presence of H3K27me3 and H3K9me3 modifications involved in both facultative and constitutive heterochromatin formation respectively.

The lower accuracy in predicting TAD boundaries may indicate that other factors in addition to histone modifications and CTCF may play a role in specification of these boundary regions. The principles of TAD formation are an ongoing field of research and it is believed that there are different classes of TADs with different functionalities. As we and others have shown, a high proportion of TAD boundaries (65-90% depending on cell type) are associated with CTCF [24]. However, there is also a significant fraction of TAD boundaries that are CTCF independent and are unaffected by CTCF loss [95]. It is also possible that demarcation of certain TAD boundaries are specified by types of histone modifications or by other genomic features that are currently not well studied. Their role in gene regulation is demonstrated when disruption of certain TAD boundaries causes ectopic chromosomal contacts and long-range transcriptional dysregulation across boundaries [22]. It is believed that these boundaries are associated with transcription [24, 96] or correspond to a division between active and repressed chromatin regions, i.e. similar to the mutually exclusive A and B chromatin compartment organization containing active and inactive domains observed across a larger genomic scale [23, 97]. There is emerging evidence that TADs (and their boundary regions) may belong to distinct functional classes. Mutations and genetic aberrations in the TAD-spanning WNT6/IHH/EPHA4/PAX3 locus are believed to cause defects in limb development. CRISPR/Cas9 genome editing of the CTCF binding sites in TAD boundary region resulted in enhancer-promoter “rewiring” leading to ectopic gene expression [98] associated with defects in limb development. Conversely, disruption of TAD boundaries through the removal of all boundary associated CTCF sites at the *Sox9-Kcnj2* locus which regulate mouse limb bud development resulted in the fusion of neighbouring TADs but had very little effects on gene expression [99]. Different domain boundary classes may have different requirements for CTCF and other factors.

In conclusion, we demonstrated that our machine and deep learning models can predict genomic elements with potential transcriptional and architectural functions usually detected by 3C methods using only histone modification and CTCF ChIP-seq data with high accuracy. We also show that the histone signatures demarcating these elements are cell type independent, where models trained on one cell type can be used to identify these elements in other cell types with comparable accuracy.

## Data Availability

All code for this project was produced in Python Jupyter or R notebooks and will be made available through GitHub after peer reviewed publication. Bash scripts, R and Python code were appropriately annotated for future usage. Only publicly available data sets were used.

## Author Contribution

S.A.S conceived the study. L.D.M carried out the data processing, genomic data analysis and developed the machine learning models. O.F contributed to the initial machine learning modelling and analysis during a summer studentship. L.P and D.B were involved with data processing and analysis. L.D.M and S.A.S wrote the manuscript with input from the other authors.

## Acknowledgements

This work was supported by UK Medical Research Council core funding (MC UU 12022/10) to S.A.S. L.P is supported by a Medical Research Council Doctoral Training Award. L.D.M., O.P, were supported by MRC core funding. We also thank members of the S.A.S laboratory that read and commented on the manuscript.

## Supplementary Data

### Figures

**Fig. S1:**
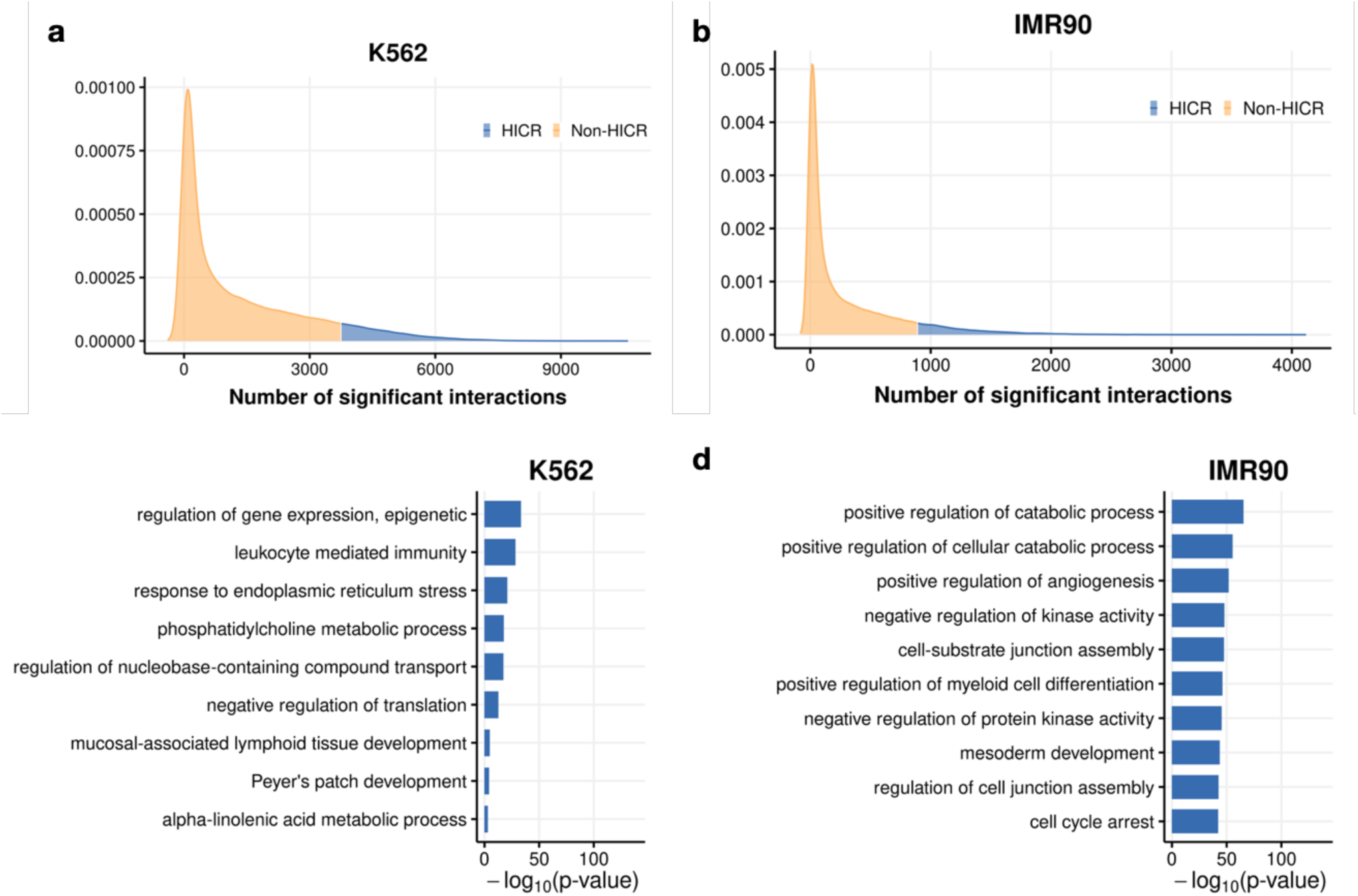
Distribution of significant interaction counts and GREAT analysis. **a-b**, The distribution of significant interaction counts for the K562 (**a**) and IMR90 (**b**) cell line. Shaded in blue are the top 10% of all interactions, which were defined as Highly Interacting Chromatin Regions (HICRs). **c-d**, Functional enrichment analysis for HICRs in the K562 (**c**) and IMR90 (**d**) cell lines. The top 10 terms are displayed, ordered by the negative log10 of P-value, which is displayed on the x-axis.

**Fig. S2:**
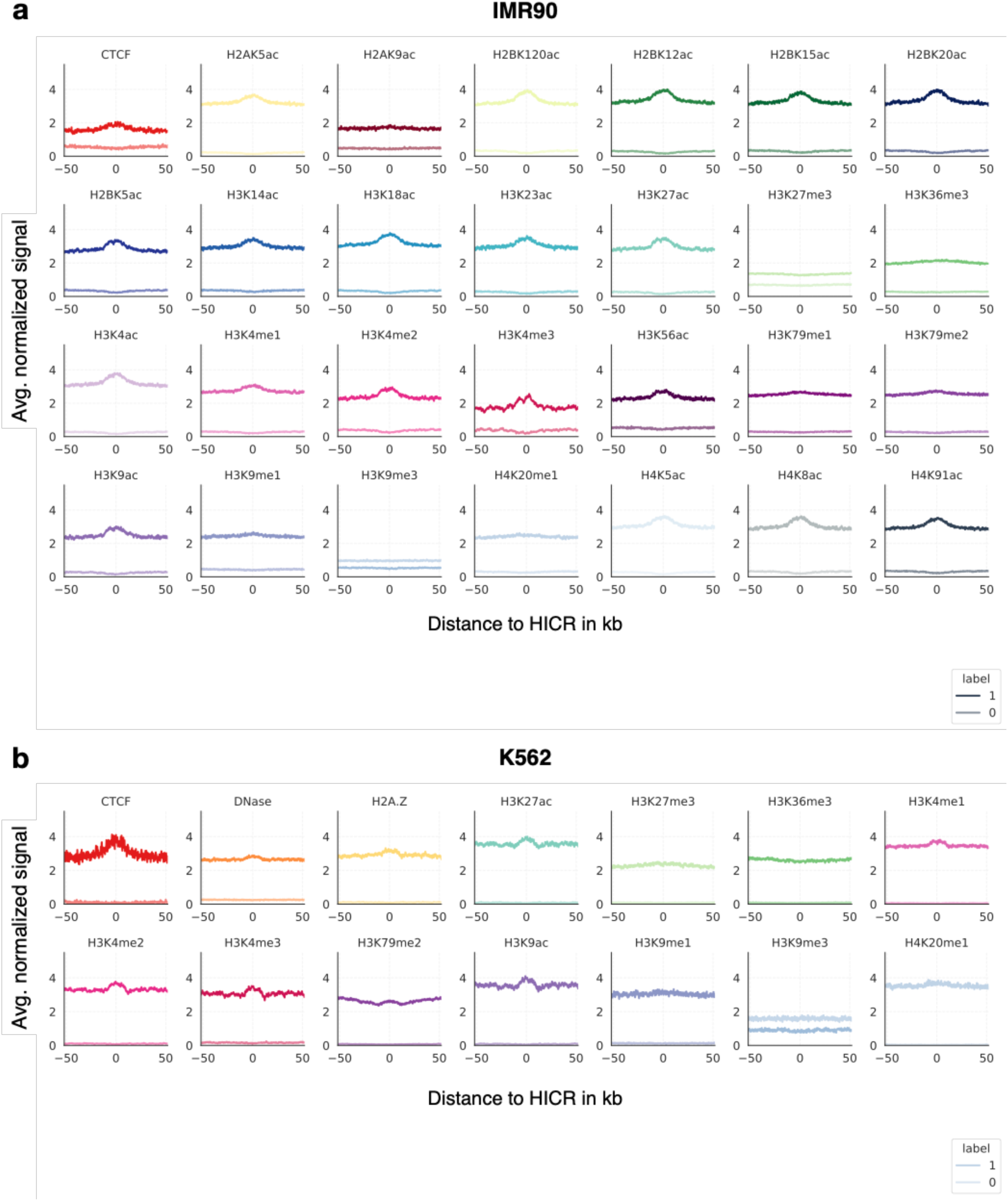
Histone mark signal at HICRs. Spatial pattern of histone modification distribution found in a 100 Kb window encapsulating the HICRs for the IMR90 (**a**) and K562 (**b**) cell line. The signal was averaged over all samples. The average signal over non-HICRs is illustrated in a lighter colour.

**Fig. S3:**
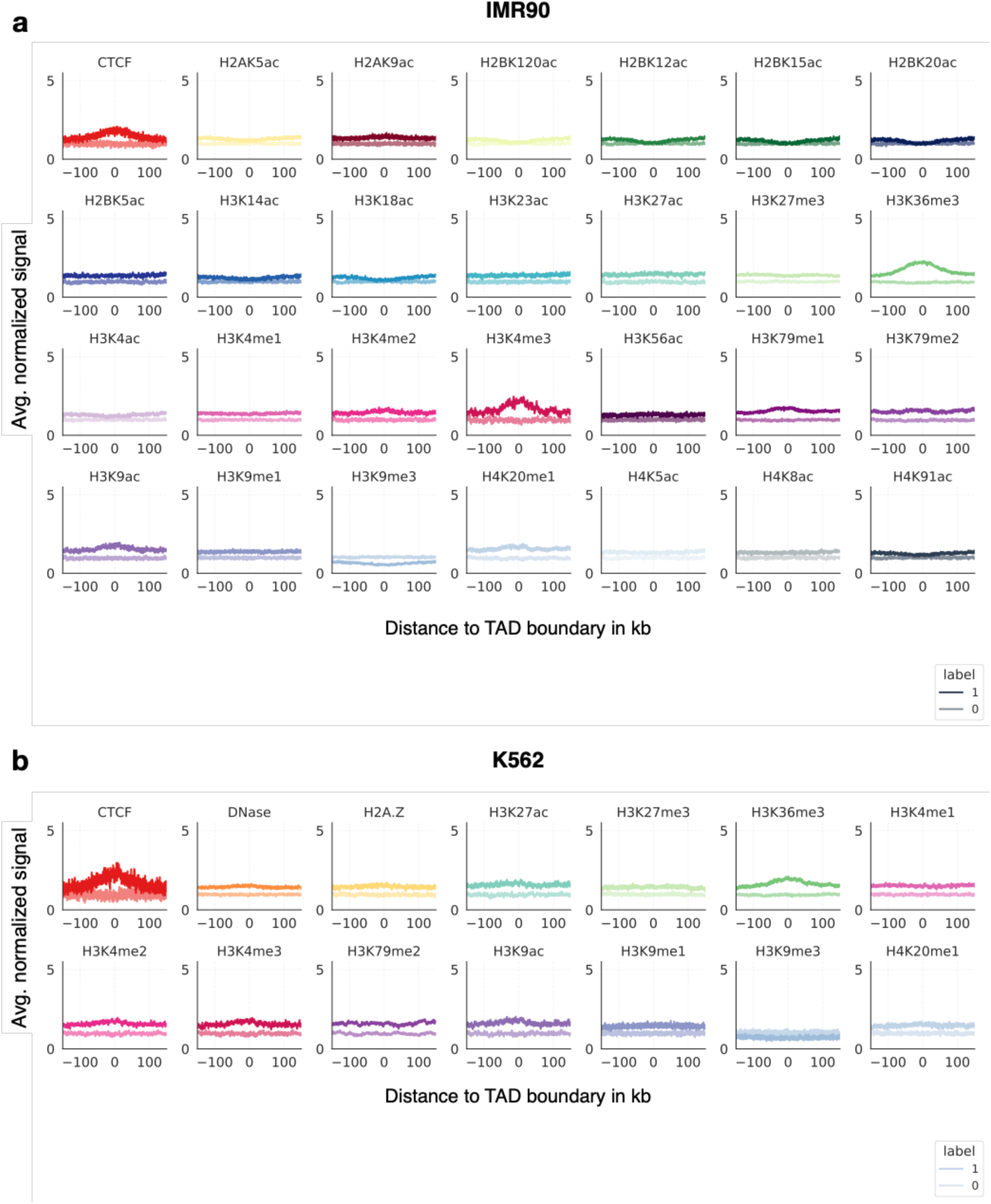
Histone mark signal at TAD boundaries. Spatial pattern of the histone modification distribution found in a 300 Kb window encapsulating the TAD boundaries for the IMR90 (**a**) and K562 (**b**) cell line. The signal was averaged over all samples. The average signal over non-TAD boundaries is illustrated in a lighter colour.

**Fig. S4:**
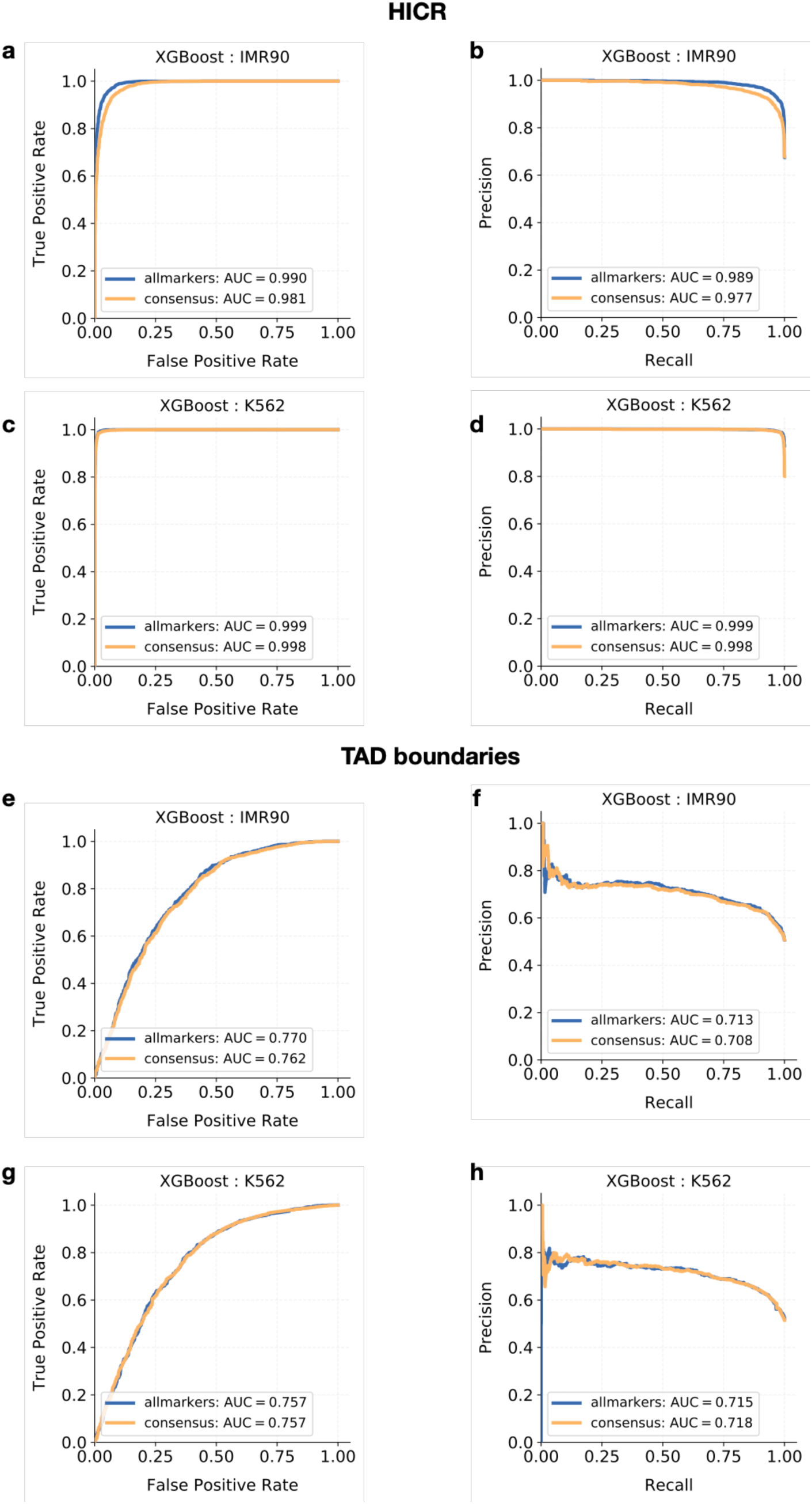
XGBoost performance on IMR90 and K562. **a,c,e,g** Receiver-operating characteristic curve and **b,d,f,h** precision-recall curve for the binary classification task of predicting HICRs (**a-d**) and TAD boundaries (**e-h**) from histone modification data. Models were trained once using all available histone modifications for each cell-type (blue) and using only the consensus set (orange).

**Fig. S5:**
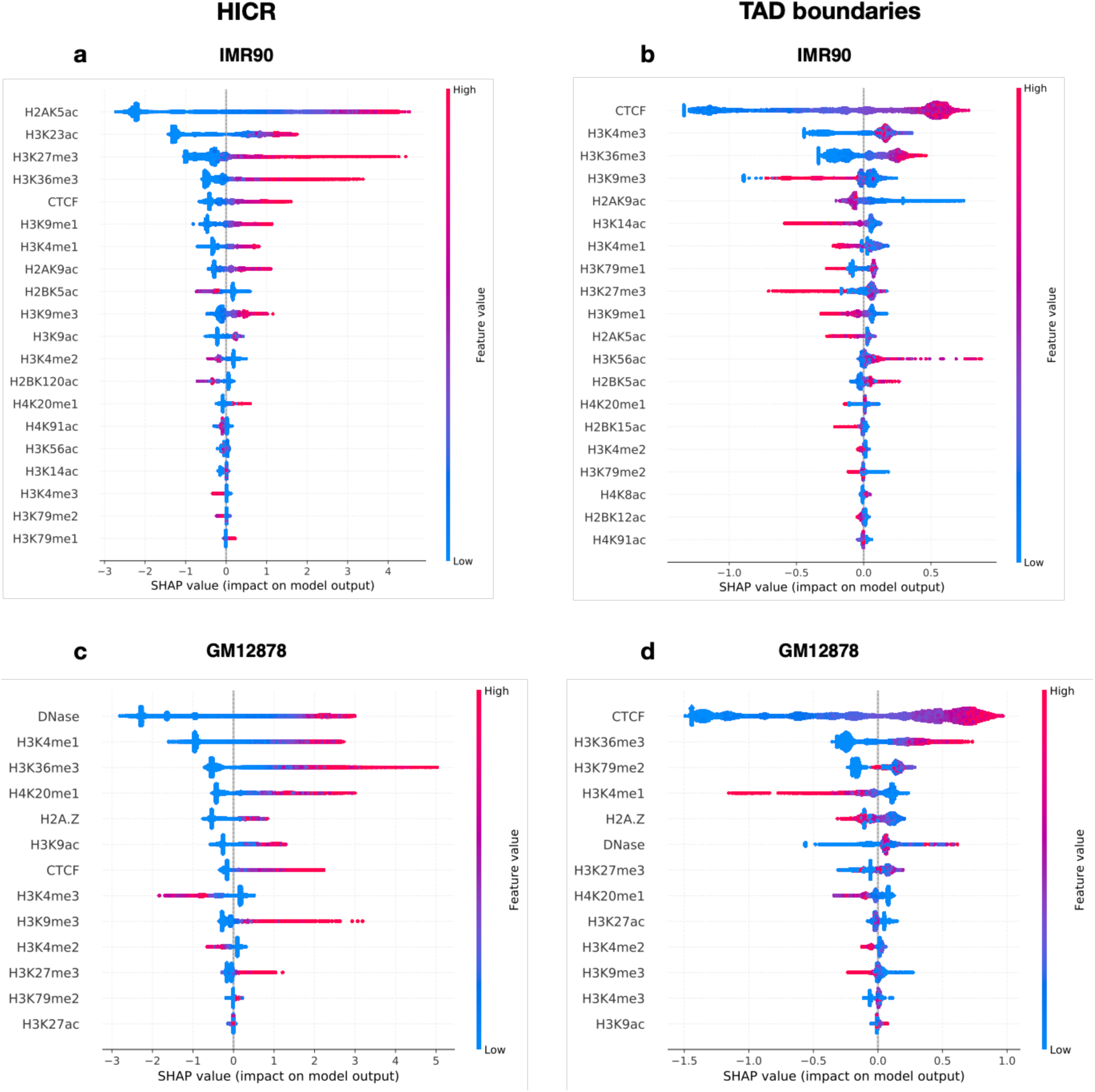
Feature importance for IMR90 and K562. Histone marks ordered by mean absolute SHAP values in HICRs (**a,c**) and TAD boundaries (**b,d**). On the y axis, the violin plot shows the full distribution of the SHAP values for each feature. The dot plot in the foreground shows a color coding of the actual value of the feature, resulting in the SHAP value as indicated on the x axis.

**Fig. S6:**
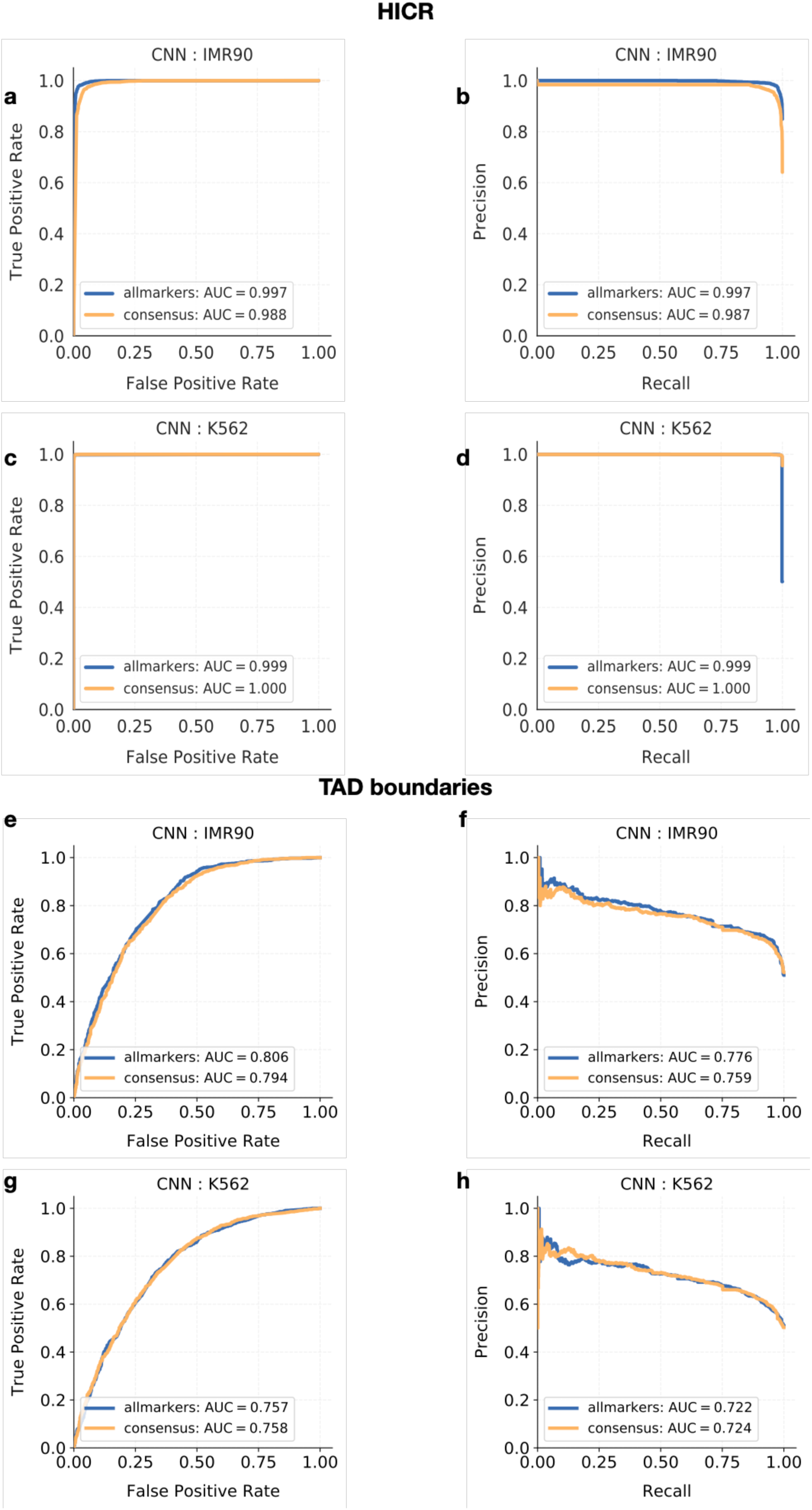
CNN performance on IMR90 and K562. **a,c,e,g** Receiver-operating characteristic curve and **b,d,f,h** precision-recall curve for the binary classification task of predicting HICRs (**a-d**) and TAD boundaries (**e-h**) from histone modification data. Models were trained once using all available histone marks for each cell-type (blue) and using only the consensus set (orange).

**Fig. S7:**
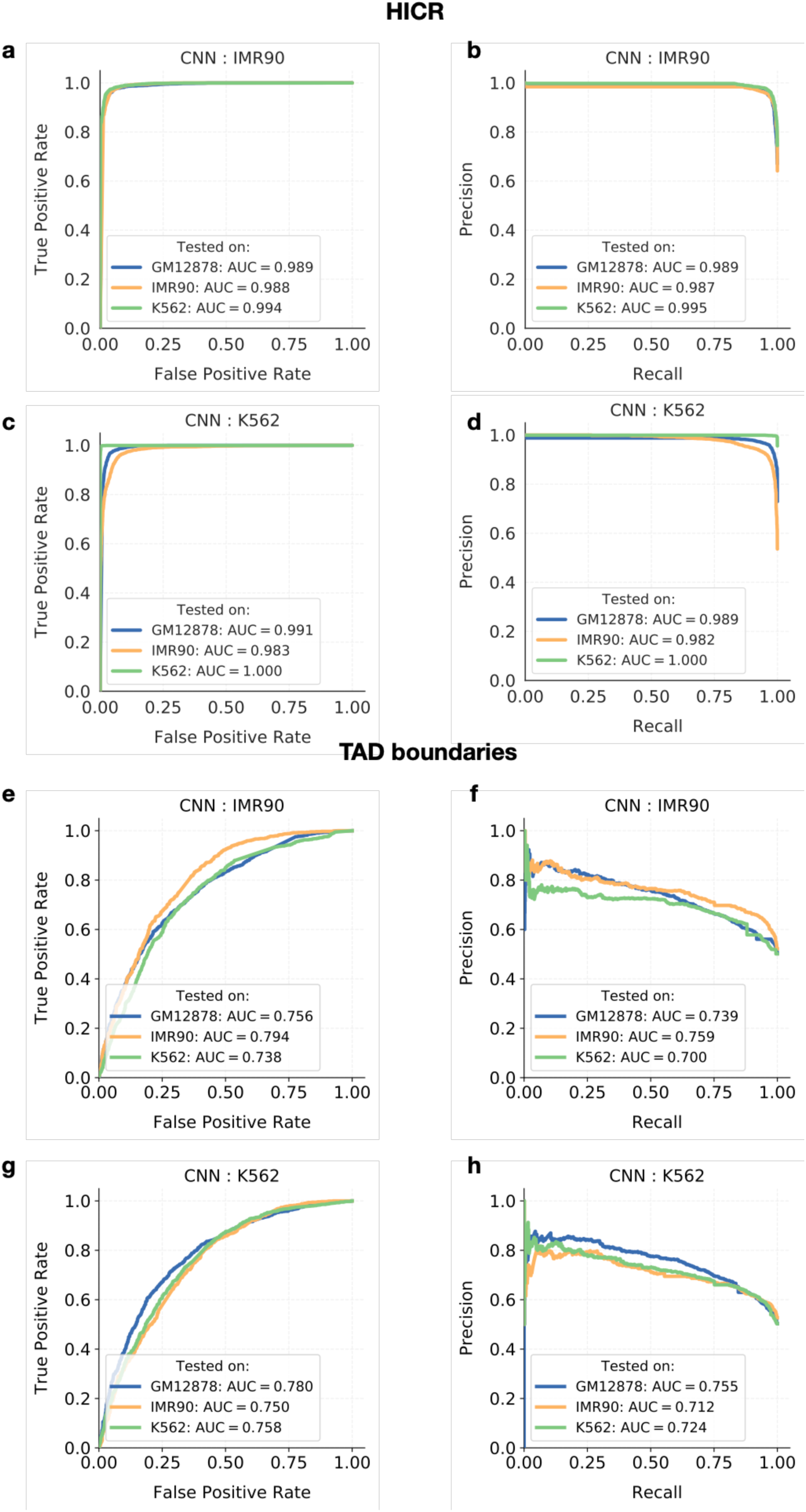
Model generalisability on IMR90 and K562 cell-types. **a,c,e,g** Receiver-operating characteristic curve and **b,d,f,h** precision-recall curve for the binary classification task of predicting HICRs (**a-d**) and TAD boundaries (**e-h**) in all three cell-types from models solely trained on the respective cell-type data. Models were trained using only the consensus set of histone modifications.

**Fig. S8:**
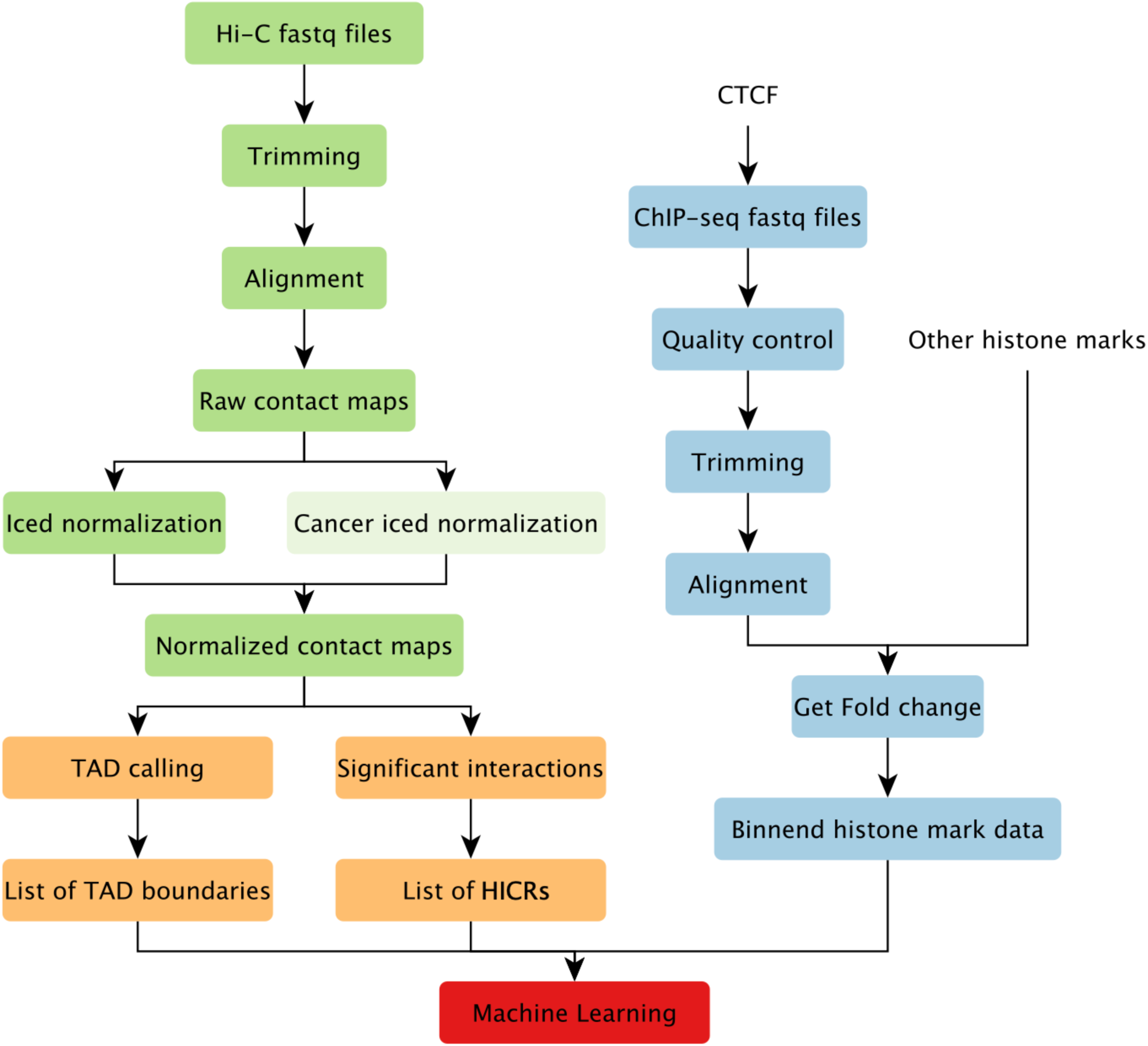
Data pre-processing. In green: Hi-C paired end reads were downloaded as fastq files. After an initial trimming step, they were aligned to the reference genome to create the raw interaction matrix. TAD calling and identification of significant interactions was performed on the normalised contact map. A list of TAD boundaries and HICRs functioned as labels for downstream supervised machine learning. In blue: If processed data was not available (CTCF), ChIP-seq fastq files were downloaded. After an initial quality control and trimming step, they were aligned to the reference genome to get fold-change values. The fold-change values were aggregated to create binned features as input for the machine learning approaches.

**Fig. S9:**
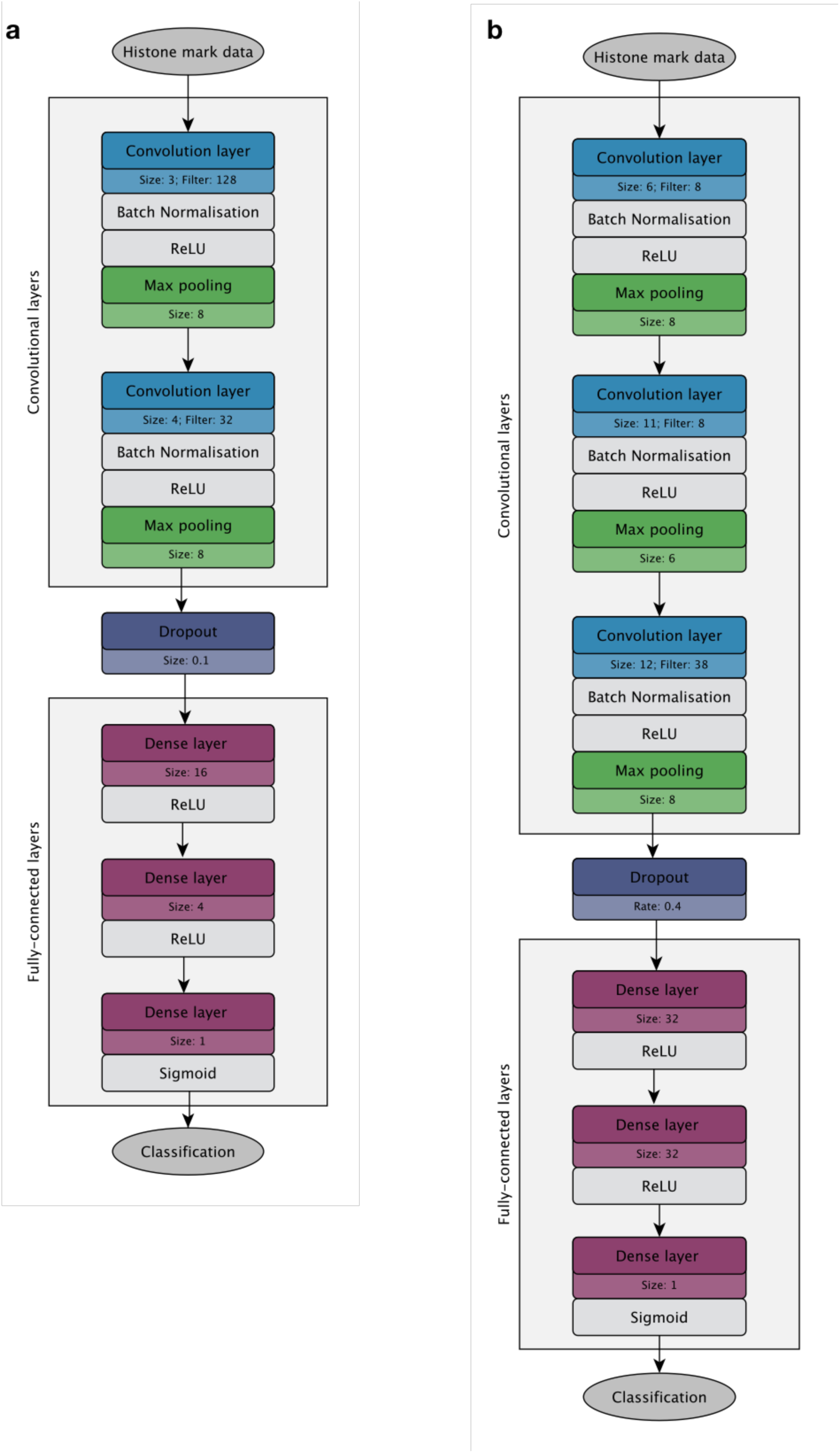
Architecture of the convolutional neural networks. If parameters differed from the default settings, they are indicated in the box below the layer. **a**, The CNN for HICRs consists of two convolution blocks followed by a dropout layer and two hidden fully connected layers. **b**, The CNN for TAD boundaries consists of three convolution blocks followed by a dropout layer and two hidden fully connected layers.

### Tables

**Table S1:** Cell type, accession number and number of reads of all CTCF ChIP-seq data sets used.

| cell type | Data-set | Accession number | Number of reads |
| --- | --- | --- | --- |
| IMR90 | Replicate 1.1 | SRR639078 | 8,913,904 |
|  | Replicate 1.2 | SRR639079 | 12,556,107 |
| GM12878 | Replicate 1.1 | SRR227594 | 29,883,393 |
|  | Replicate 1.2 | SRR227595 | 17,949,455 |
|  | Input | SRR299277 | 16,990,693 |
|  | Replicate 2 | SRR299312 | 14,311,175 |
|  | Replicate 3 | SRR299314 | 31,495,220 |
|  | Replicate 4.1 | SRR357519 | 8,420,728 |
|  | Replicate 4.2 | SRR357520 | 16,281,148 |
|  | Replicate 5 | SRR502628 | 29,485,157 |
|  | Replicate 5 | SRR502629 | 16,411,438 |
|  | Replicate 4.3 | SRR585830 | 21,319,387 |
| K562 | Replicate 1.1 | SRR227523 | 19,656,110 |
|  | Replicate 1.2 | SRR227524 | 16,448,282 |
|  | Replicate 2 | SRR299298 | 32,740,518 |
|  | Replicate 3 | SRR299327 | 18,123,856 |
|  | Replicate 4 | SRR299341 | 27,953,212 |
|  | Replicate 5.1 | SRR577849 | 24,638,860 |
|  | Replicate 5.2 | SRR577850 | 31,332,287 |

**Table S2:** Cell type, accession number and number of reads of all Hi-C data-sets used. All runs from the experiments were used.

| cell type | Data-set | Accession number | Number of reads |
| --- | --- | --- | --- |
| IMR90 | Replicate 1 | SRX212172 | 397,194,480 |
|  | Replicate 2 | SRX212173 | 440,242,130 |
|  | Replicate 3 | SRX294948 | 621,089,009 |
|  | Replicate 4 | SRX294949 | 529,157,703 |
|  | Replicate 5 | SRX294950 | 234,133,577 |
|  | Replicate 6 | SRX294951 | 208,710,657 |
| GM12878 | Replicate 1 | SRX764940 | 304,049,399 |
|  | Replicate 2 | SRX764958 | 322,202,135 |
|  | Replicate 3 | SRX764960 | 347,956,701 |
| K562 | Replicate 1 | SRX765004 | 456,757,799 |
|  | Replicate 2 | SRX765005 | 591,854,553 |
|  | Replicate 3 | SRX765006 | 79,905,895 |
|  | Replicate 4 | SRX765007 | 79,578,049 |

**Table S3:** Genomic Association Test (GAT) analysis of HICRs to identify significance of overlap with multiple sets of genomic intervals. Displayed are the number of overlapping segments, the expected number of overlapping segments from 3000 permutations, the fold change of observed to expected overlap and a multi-testing corrected q-value.

| Database | cell type | observed overlap | expected overlap | fold change | q-value |
| --- | --- | --- | --- | --- | --- |
| Conservation UCSC | GM12878 | 13708 | 11834 | 1.16 | 0.0003 |
|  | IMR90 | 12014 | 10690 | 1.12 | 0.0003 |
|  | K562 | 13103 | 11094 | 1.18 | 0.0003 |
| CpG Islands | GM12878 | 4785 | 2232 | 2.14 | 0.0003 |
|  | IMR90 | 2259 | 1599 | 1.41 | 0.0003 |
|  | K562 | 5351 | 2260 | 2.37 | 0.0003 |
| dbSUPER | GM12878 | 637 | 80 | 7.86 | 0.0003 |
|  | IMR90 | 518 | 98 | 5.26 | 0.0003 |
|  | K562 | 729 | 196 | 3.71 | 0.0003 |
| EnhancerAtlas 2.0 | GM12878 | 9794 | 3522 | 2.78 | 0.0003 |
|  | IMR90 | 9062 | 4247 | 2.13 | 0.0003 |
|  | K562 | 8753 | 3172 | 2.76 | 0.0003 |
| Fantom5 Enhancers | GM12878 | 7398 | 3804 | 1.94 | 0.0004 |
|  | IMR90 | 5986 | 3214 | 1.86 | 0.0004 |
|  | K562 | 6436 | 3693 | 1.74 | 0.0004 |
| Ensembl Genes | GM12878 | 10866 | 6739 | 1.61 | 0.0003 |
|  | IMR90 | 8040 | 5724 | 1.40 | 0.0003 |
|  | K562 | 9909 | 6461 | 1.53 | 0.0003 |
| Ensembl Regulatory Build | GM12878 | 10483 | 5153 | 2.03 | 0.0004 |
|  | IMR90 | 7299 | 4181 | 1.75 | 0.0004 |
|  | K562 | 10157 | 5002 | 2.03 | 0.0004 |
| GeneHancer | GM12878 | 13707 | 10919 | 1.26 | 0.0003 |
|  | IMR90 | 11990 | 9722 | 1.23 | 0.0003 |
|  | K562 | 13100 | 10322 | 1.27 | 0.0003 |
| HumCFS Fragile Sites | GM12878 | 5117 | 3728 | 1.37 | 0.0003 |
|  | IMR90 | 3998 | 3258 | 1.23 | 0.0003 |
|  | K562 | 5622 | 3695 | 1.52 | 0.0003 |
| Microsatellites UCSC | GM12878 | 3100 | 3000 | 1.03 | 0.0080 |
|  | IMR90 | 3063 | 2689 | 1.14 | 0.0006 |
|  | K562 | 2950 | 2852 | 1.03 | 0.0142 |
| Repeatmasker UCSC | GM12878 | 13708 | 11865 | 1.16 | 0.0003 |
|  | IMR90 | 12014 | 10713 | 1.12 | 0.0003 |
|  | K562 | 13103 | 11133 | 1.18 | 0.0003 |
| SEdb:Enhancer_Element | GM12878 | 446 | 53 | 8.25 | 0.0004 |
|  | IMR90 | 601 | 142 | 4.22 | 0.0003 |
|  | K562 | 684 | 182 | 3.75 | 0.0004 |
| SEdb:Super_Enhancer | GM12878 | 446 | 54 | 8.12 | 0.0004 |
|  | IMR90 | 601 | 141 | 4.23 | 0.0003 |
|  | K562 | 684 | 181 | 3.75 | 0.0004 |
| SEdb:Typical_Enhancer | GM12878 | 5842 | 1465 | 3.98 | 0.0003 |
|  | IMR90 | 4153 | 1330 | 3.12 | 0.0003 |
|  | K562 | 3841 | 1400 | 2.74 | 0.0003 |
| TF clusters UCSC | GM12878 | 13702 | 11711 | 1.17 | 0.0003 |
|  | IMR90 | 12012 | 10569 | 1.14 | 0.0003 |
|  | K562 | 13093 | 10981 | 1.19 | 0.0003 |
| TSS Ensembl | GM12878 | 7438 | 3566 | 2.09 | 0.0004 |
|  | IMR90 | 4334 | 2745 | 1.58 | 0.0004 |
|  | K562 | 7215 | 3529 | 2.04 | 0.0004 |
| TSS Fantom | GM12878 | 13155 | 9131 | 1.44 | 0.0003 |
|  | IMR90 | 10969 | 8011 | 1.37 | 0.0003 |
|  | K562 | 12369 | 8667 | 1.43 | 0.0003 |

**Table S4:** Genomic Association Test (GAT) analysis of TAD boundaries to identify significance of overlap with multiple sets of genomic intervals. Displayed are the number of overlapping segments, the expected number of overlapping segments from 3000 permutations, the fold change of observed to expected overlap and a multi-testing corrected q-value.

| Database | cell type | observed overlap | expected overlap | fold change | q-value |
| --- | --- | --- | --- | --- | --- |
| Conservation UCSC | GM12878 | 6224 | 5685 | 1.09 | 0.0003 |
|  | IMR90 | 5368 | 4942 | 1.09 | 0.0003 |
|  | K562 | 6275 | 5685 | 1.10 | 0.0003 |
| CpG Islands | GM12878 | 2505 | 1423 | 1.76 | 0.0003 |
|  | IMR90 | 2398 | 1210 | 1.98 | 0.0003 |
|  | K562 | 2505 | 1419 | 1.76 | 0.0003 |
| dbSUPER | GM12878 | 51 | 43 | 1.17 | 0.1337 |
|  | IMR90 | 77 | 57 | 1.34 | 0.0003 |
|  | K562 | 159 | 95 | 1.66 | 0.0003 |
| EnhancerAtlas 2.0 | GM12878 | 3086 | 2177 | 1.42 | 0.0003 |
|  | IMR90 | 3544 | 2574 | 1.38 | 0.0003 |
|  | K562 | 2955 | 1825 | 1.62 | 0.0003 |
| Fantom5 Enhancers | GM12878 | 3011 | 2555 | 1.18 | 0.0003 |
|  | IMR90 | 2678 | 2219 | 1.21 | 0.0003 |
|  | K562 | 3192 | 2559 | 1.25 | 0.0003 |
| Ensembl Genes | GM12878 | 4516 | 3290 | 1.37 | 0.0003 |
|  | IMR90 | 4130 | 2860 | 1.44 | 0.0003 |
|  | K562 | 4527 | 3285 | 1.38 | 0.0003 |
| Ensembl Regulatory Build | GM12878 | 4180 | 3101 | 1.35 | 0.0003 |
|  | IMR90 | 3813 | 2687 | 1.42 | 0.0003 |
|  | K562 | 4290 | 3088 | 1.39 | 0.0003 |
| GeneHancer | GM12878 | 5998 | 5184 | 1.16 | 0.0003 |
|  | IMR90 | 5227 | 4510 | 1.16 | 0.0003 |
|  | K562 | 6076 | 5165 | 1.18 | 0.0003 |
| HumCFS Fragile Sites | GM12878 | 1856 | 1669 | 1.11 | 0.0003 |
|  | IMR90 | 1673 | 1486 | 1.13 | 0.0003 |
|  | K562 | 1861 | 1634 | 1.14 | 0.0003 |
| Microsatellites UCSC | GM12878 | 2510 | 2469 | 1.02 | 0.2575 |
|  | IMR90 | 2151 | 2148 | 1.00 | 0.4300 |
|  | K562 | 2498 | 2465 | 1.01 | 0.2575 |
| Repeatmasker UCSC | GM12878 | 6270 | 5699 | 1.10 | 0.0003 |
|  | IMR90 | 5404 | 4954 | 1.09 | 0.0003 |
|  | K562 | 6322 | 5699 | 1.11 | 0.0003 |
| SEdb: Enhancer_Element | GM12878 | 36 | 27 | 1.31 | 0.0600 |
|  | IMR90 | 123 | 84 | 1.45 | 0.0003 |
|  | K562 | 142 | 87 | 1.62 | 0.0003 |
| SEdb: Super_Enhancer | GM12878 | 36 | 28 | 1.28 | 0.1000 |
|  | IMR90 | 123 | 84 | 1.45 | 0.0003 |
|  | K562 | 142 | 88 | 1.61 | 0.0003 |
| SEdb: Typical_Enhancer | GM12878 | 1282 | 954 | 1.34 | 0.0003 |
|  | IMR90 | 1256 | 962 | 1.31 | 0.0003 |
|  | K562 | 1338 | 925 | 1.45 | 0.0003 |
| TF clusters UCSC | GM12878 | 6202 | 5646 | 1.10 | 0.0003 |
|  | IMR90 | 5339 | 4924 | 1.08 | 0.0003 |
|  | K562 | 6197 | 5633 | 1.10 | 0.0003 |
| TSS Ensembl | GM12878 | 3476 | 2148 | 1.62 | 0.0003 |
|  | IMR90 | 3339 | 1860 | 1.79 | 0.0003 |
|  | K562 | 3554 | 2142 | 1.66 | 0.0003 |
| TSS Fantom | GM12878 | 5718 | 4939 | 1.16 | 0.0003 |
|  | IMR90 | 5014 | 4300 | 1.17 | 0.0003 |
|  | K562 | 5793 | 4928 | 1.18 | 0.0003 |

**Table S5:** List of available ChIP-seq data-sets for each cell-type and the consensus set.

|  | IMR90 | GM12878 | K562 | consensus |
| --- | --- | --- | --- | --- |
| H2AK5ac | ✓ |  |  |  |
| H2AK9ac | ✓ |  |  |  |
| H2BK5ac | ✓ |  |  |  |
| H2BK12ac | ✓ |  |  |  |
| H2BK15ac | ✓ |  |  |  |
| H2BK20ac | ✓ |  |  |  |
| H2BK120ac | ✓ |  |  |  |
| H3K4ac | ✓ |  |  |  |
| H3K4me1 | ✓ | ✓ | ✓ | ✓ |
| H3K4me2 | ✓ | ✓ | ✓ | ✓ |
| H3K4me3 | ✓ | ✓ | ✓ | ✓ |
| H3K9ac | ✓ | ✓ | ✓ | ✓ |
| H3K9me1 | ✓ |  | ✓ |  |
| H3K9me3 | ✓ | ✓ | ✓ | ✓ |
| H3K14ac | ✓ |  |  |  |
| H3K18ac | ✓ |  |  |  |
| H3K23ac | ✓ |  |  |  |
| H3K27ac | ✓ | ✓ | ✓ | ✓ |
| H3K27me3 | ✓ | ✓ | ✓ | ✓ |
| H3K36me3 | ✓ | ✓ | ✓ | ✓ |
| H3K56ac | ✓ |  |  |  |
| H3K79me1 | ✓ |  |  |  |
| H3K79me2 | ✓ | ✓ | ✓ | ✓ |
| H4K5ac | ✓ |  |  |  |
| H4K8ac | ✓ |  |  |  |
| H4K20me1 | ✓ | ✓ | ✓ | ✓ |
| H4K91ac | ✓ |  |  |  |
| CTCF | ✓ | ✓ | ✓ | ✓ |
| DHS assay |  | ✓ | ✓ |  |
| H2A.Z |  | ✓ | ✓ |  |

**Table S6:** Logistic Regression -Hyperparameter space and optimal parameters

| parameter | space | optimal parameter |  |
| --- | --- | --- | --- |
|  |  | HICR | TAD boundary |
| penalty | [none , L2] | L2 | L2 |
| C | loguniform: 0.001 – 10 | 3.762 | 0.460 |
| solver | ['lbfgs', 'saga'] | 'saga' | 'lbfgs' |
| max_iter | [100, 150, 200, 250, 300] | 100 | 100 |

**Table S7:** XGBoost: Hyperparameter space and optimal parameters

| parameter | space | optimal parameter |  |
| --- | --- | --- | --- |
|  |  | HICR | TAD boundary |
| max_depth | [3, 2] | 3 | 3 |
| n_estimator | 300, 310, 320, ..., 1200 | 560 | 870 |
| subsample | [0.75, 0.5, 0.25] | 0.5 | 0.75 |
| colsample_bytree | [0.75, 0.5, 0.25, 0.125] | 0.75 | 0.75 |
| learning_rate | loguniform: 0.00001 - 0.001 | 0.0496 | 0.0496 |

**Table S8:**
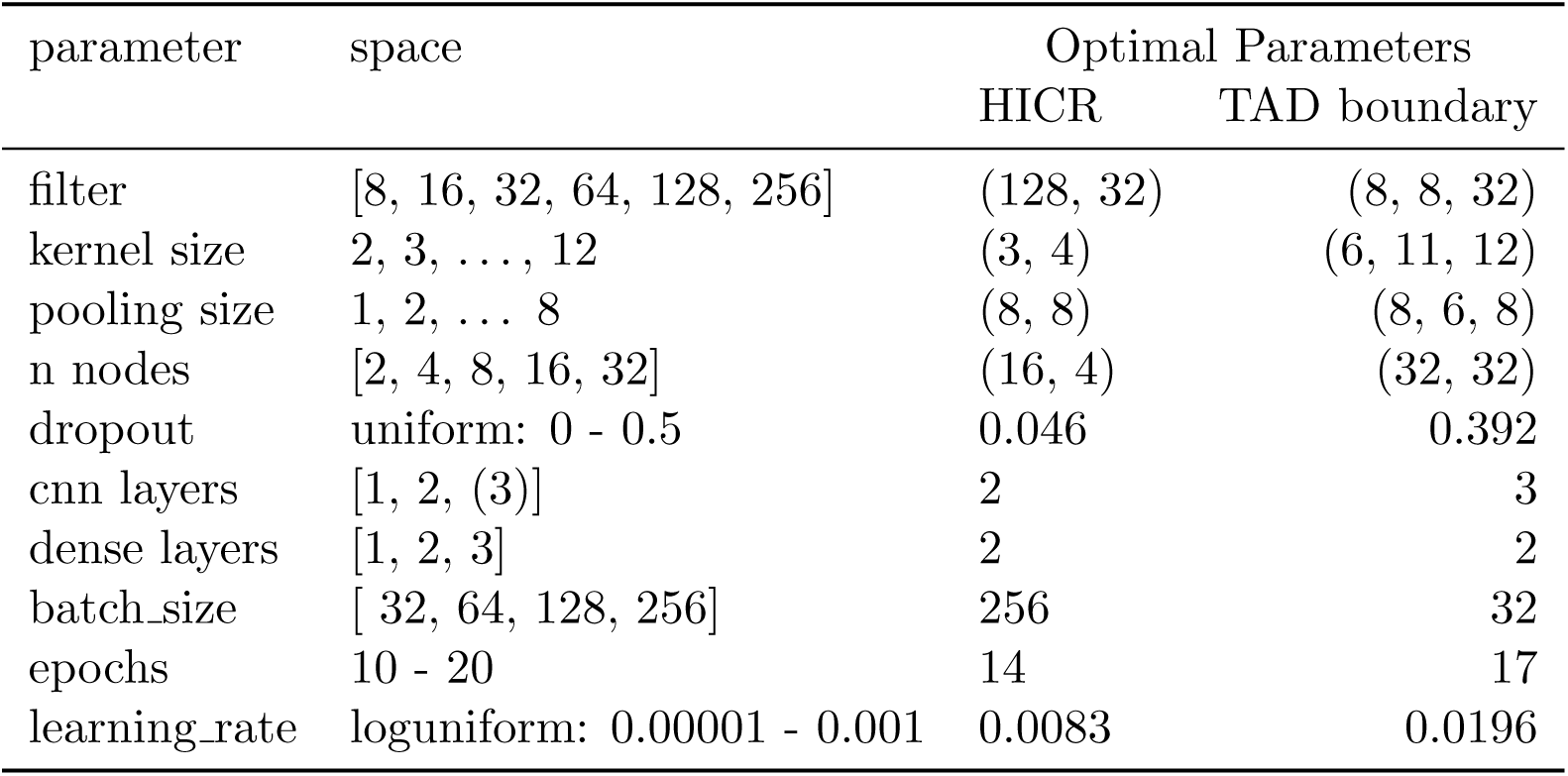
CNN: Hyperparameter space and optimal parameters

| parameter | space | Optimal Parameters |  |
| --- | --- | --- | --- |
|  |  | HICR | TAD boundary |
| filter | [8, 16, 32, 64, 128, 256] | (128, 32) | (8, 8, 32) |
| kernel size | 2, 3, ..., 12 | (3, 4) | (6, 11, 12) |
| pooling size | 1, 2, ..., 8 | (8, 8) | (8, 6, 8) |
| n nodes | [2, 4, 8, 16, 32] | (16, 4) | (32, 32) |
| dropout | uniform: 0 - 0.5 | 0.046 | 0.392 |
| cnn layers | [1, 2, (3)] | 2 | 3 |
| dense layers | [1, 2, 3] | 2 | 2 |
| batch_size | [ 32, 64, 128, 256] | 256 | 32 |
| epochs | 10 - 20 | 14 | 17 |
| learning_rate | loguniform: 0.00001 - 0.001 | 0.0083 | 0.0196 |

**Table S9:** HICRs: Different metrics for evaluation of Logistic Regression model.

| celltype | markers | Acc. | Recall | Precision | F1 | ROC AUC | PR AUC |
| --- | --- | --- | --- | --- | --- | --- | --- |
| GM12878 | allmarkers | 0.951 | 0.930 | 0.966 | 0.947 | 0.990 | 0.990 |
|  | consensus | 0.946 | 0.920 | 0.964 | 0.942 | 0.987 | 0.987 |
| IMR90 | allmarkers | 0.951 | 0.947 | 0.947 | 0.947 | 0.990 | 0.988 |
|  | consensus | 0.925 | 0.907 | 0.931 | 0.919 | 0.979 | 0.976 |
| K562 | allmarkers | 0.984 | 0.981 | 0.988 | 0.984 | 0.999 | 0.998 |
|  | consensus | 0.981 | 0.975 | 0.988 | 0.981 | 0.998 | 0.997 |

**Table S10:** TAD boundaries: Different metrics for evaluation of Logistic Regression model.

| celltype | markers | Acc. | Recall | Precision | F1 | ROC AUC | PR AUC |
| --- | --- | --- | --- | --- | --- | --- | --- |
| GM12878 | allmarkers | 0.689 | 0.670 | 0.685 | 0.678 | 0.754 | 0.686 |
|  | consensus | 0.687 | 0.663 | 0.685 | 0.674 | 0.750 | 0.683 |
| IMR90 | allmarkers | 0.704 | 0.717 | 0.691 | 0.704 | 0.764 | 0.698 |
|  | consensus | 0.699 | 0.703 | 0.690 | 0.696 | 0.754 | 0.690 |
| K562 | allmarkers | 0.681 | 0.668 | 0.697 | 0.682 | 0.740 | 0.705 |
|  | consensus | 0.677 | 0.662 | 0.693 | 0.677 | 0.738 | 0.704 |

## References

[1] T. Cremer and C. Cremer, “Chromosome territories, nuclear architecture and gene regulation in mammalian cells,” Nature Reviews Genetics, vol. 2, no. 4, pp. 292–301, 2001.

[2] T. Sexton, H. Schober, P. Fraser, and S. M. Gasser, “Gene regulation through nuclear organization,” Nature Structural and Molecular Biology, vol. 14, no. 11, pp. 1049–1055, 2007.

[3] W. A. Bickmore, “The Spatial Organization of the Human Genome,” Annual Review of Genomics and Human Genetics, vol. 14, no. 1, pp. 67–84, 2013.

[4] K. Luger, A. W. Mäder, R. K. Richmond, D. F. Sargent, and T. J. Richmond, “Crystal structure of the nucleosome core particle at 2.8 åresolution,” Nature, vol. 389, no. 6648, pp. 251–260, 1997.

[5] V. Allfrey, R. Faulkner, and A. Mirsky, “Acetylation and methylation of histones and their possible role in the regulation of rna synthesis,” Proceedings of the National Academy of Sciences, vol. 51, no. 5, pp. 786–794, 1964.

[6] A. J. Ruthenburg, H. Li, D. J. Patel, and C. D. Allis, “Multivalent engagement of chromatin modifications by linked binding modules,” Nature reviews Molecular cell biology, vol. 8, no. 12, pp. 983–994, 2007.

[7] C. D. Allis, L. M. Caparros, T. Jenuwein, and D. Reinberg, Epigenetics. Cold Spring Harbor, NY: Cold Spring Harbor Laboratory Press, 2nd ed., 2015.

[8] J. R. Ecker, W. A. Bickmore, I. Barroso, J. K. Pritchard, Y. Gilad, and E. Segal, “ENCODE explained,” Nature, vol. 489, p. 52, sep 2012.

[9] T. J. Stevens, D. Lando, S. Basu, L. P. Atkinson, Y. Cao, S. F. Lee, M. Leeb, K. J. Wohlfahrt, W. Boucher, A. O’Shaughnessy-Kirwan, et al., “3d structures of individual mammalian genomes studied by single-cell hi-c,” Nature, vol. 544, no. 7648, pp. 59–64, 2017.

[10] G. Li, X. Ruan, R. Auerbach, K. Sandhu, M. Zheng, P. Wang, H. Poh, Y. Goh, J. Lim, J. Zhang, H. Sim, S. Peh, F. Mulawadi, C. Ong, Y. Orlov, S. Hong, Z. Zhang, S. Landt, D. Raha, G. Euskirchen, C.-L. Wei, W. Ge, H. Wang, C. Davis, K. I. Fisher-Aylor, A. Mortazavi, M. Gerstein, T. Gingeras, B. Wold, Y. Sun, M. Fullwood, E. Cheung, E. Liu, W.-K. Sung, M. Snyder, and Y. Ruan, “Extensive Promoter-Centered Chromatin Interactions Provide a Topological Basis for Transcription Regulation,” Cell, vol. 148, pp. 84–98, jan 2012.

[11] A. Sanyal, B. R. Lajoie, G. Jain, and J. Dekker, “The long-range interaction landscape of gene promoters,” Nature, vol. 489, pp. 109–113, sep 2012.

[12] F. Jin, Y. Li, J. R. Dixon, S. Selvaraj, Z. Ye, A. Y. Lee, C. A. Yen, A. D. Schmitt, C. A. Espinoza, and B. Ren, “A high-resolution map of the three-dimensional chromatin interactome in human cells,” Nature, vol. 503, no. 7475, pp. 290–294, 2013.

[13] J. Dowen, Z. Fan, D. Hnisz, G. Ren, B. Abraham, L. Zhang, A. Weintraub, J. Schuijers, T. Lee, K. Zhao, and R. Young, “Control of Cell Identity Genes Occurs in Insulated Neighborhoods in Mammalian Chromosomes,” Cell, vol. 159, pp. 374–387, oct 2014.

[14] N. Heidari, D. H. Phanstiel, C. He, F. Grubert, F. Jahanbani, M. Kasowski, M. Q. Zhang, and M. P. Snyder, “Genome-wide map of regulatory interactions in the human genome.,” Genome research, vol. 24, pp. 1905–17, ec 2014.

[15] M. Simonis, P. Klous, E. Splinter, Y. Moshkin, R. Willemsen, E. De Wit, B. Van Steensel, and W. De Laat, “Nuclear organization of active and inactive chromatin domains uncovered by chromosome conformation capture–on-chip (4c),” Nature genetics, vol. 38, no. 11, pp. 1348–1354, 2006.

[16] J. Dostie, T. A. Richmond, R. A. Arnaout, R. R. Selzer, W. L. Lee, T. A. Honan, E. D. Rubio, A. Krumm, J. Lamb, C. Nusbaum, et al., “Chromosome conformation capture carbon copy (5c): a massively parallel solution for mapping interactions between genomic elements,” Genome research, vol. 16, no. 10, pp. 1299–1309, 2006.

[17] E. Lieberman-Aiden, N. L. Van Berkum, L. Williams, M. Imakaev, T. Ragoczy, A. Telling, I. Amit, B. R. Lajoie, P. J. Sabo, M. O. Dorschner, et al., “Comprehensive mapping of I. long-range interactions reveals folding principles of the human genome,” science, vol. 326, no. 5950, pp. 289–293, 2009.

[18] M. J. Fullwood, M. H. Liu, Y. F. Pan, J. Liu, H. Xu, Y. B. Mohamed, Y. L. Orlov, S. Velkov, A. Ho, P. H. Mei, et al., “An oestrogen-receptor-*α*-bound human chromatin interactome,” Nature, vol. 462, no. 7269, pp. 58–64, 2009.

[19] A. D. Schmitt, M. Hu, and B. Ren, “Genome-wide mapping and analysis of chromosome architecture,” Nature Reviews Molecular Cell Biology, vol. 17, no. 12, pp. 743–755, 2016.

[20] E. Lieberman-Aiden, N. L. V. Berkum, L. Williams, M. Imakaev, T. Ragoczy, A. Telling, I. Amit, B. R. Lajoie, P. J. Sabo, M. O. Dorschner, R. Sandstrom, B. Bernstein, M. A. Bender, M. Groudine, A. Gnirke, J. Stamatoyannopoulos, and L. A. Mirny, “Comprehensive Mapping of Long-Range Interactions Reveals Folding Principles of the Human Genome,” Science, vol. 33292, no. October, pp. 289–294, 2009.

[21] J. R. Dixon, D. U. Gorkin, and B. Ren, “Chromatin Domains: The Unit of Chromosome Organization,” Molecular Cell, vol. 62, no. 5, pp. 668–680, 2016.

[22] E. P. Nora, B. R. Lajoie, E. G. Schulz, L. Giorgetti, I. Okamoto, N. Servant, T. Piolot, N. L. Van Berkum, J. Meisig, J. Sedat, J. Gribnau, E. Barillot, N. Blüthgen, J. Dekker, and E. Heard, “Spatial partitioning of the regulatory landscape of the X-inactivation centre,” Nature, vol. 485, no. 7398, pp. 381–385, 2012.

[23] S. S. Rao, M. H. Huntley, N. C. Durand, E. K. Stamenova, I. D. Bochkov, J. T. Robinson, A. L. Sanborn, I. Machol, A. D. Omer, E. S. Lander, and E. L. Aiden, “A 3D map of the human genome at kilobase resolution reveals principles of chromatin looping,” Cell, vol. 159, no. 7, pp. 1665–1680, 2014.

[24] J. R. Dixon, S. Selvaraj, F. Yue, A. Kim, Y. Li, Y. Shen, M. Hu, J. S. Liu, and B. Ren, “Topological domains in mammalian genomes identified by analysis of chromatin interactions,” Nature, vol. 485, no. 7398, pp. 376–380, 2012.

[25] B. M. Javierre, O. S. Burren, S. P. Wilder, R. Kreuzhuber, S. M. Hill, S. Sewitz, J. Cairns, S. W. Wingett, C. Várnai, M. J. Thiecke, et al., “Lineage-specific genome architecture links enhancers and non-coding disease variants to target gene promoters,” Cell, vol. 167, no. 5, pp. 1369–1384, 2016.

[26] E. Rodríguez-Carballo, L. Lopez-Delisle, Y. Zhan, P. J. Fabre, L. Beccari, I. El-Idrissi, T. H. N. Huynh, H. Ozadam, J. Dekker, and D. Duboule, “The hoxd cluster is a dynamic and resilient tad boundary controlling the segregation of antagonistic regulatory landscapes,” Genes & development, vol. 31, no. 22, pp. 2264–2281, 2017.

[27] O. Symmons, V. V. Uslu, T. Tsujimura, S. Ruf, S. Nassari, W. Schwarzer, L. Ettwiller, and F. Spitz, “Functional and topological characteristics of mammalian regulatory domains,” Genome research, vol. 24, no. 3, pp. 390–400, 2014.

[28] Y. Shen, F. Yue, D. F. McCleary, Z. Ye, L. Edsall, S. Kuan, U. Wagner, J. Dixon, L. Lee, V. V. Lobanenkov, et al., “A map of the cis-regulatory sequences in the mouse genome,” Nature, vol. 488, no. 7409, pp. 116–120, 2012.

[29] W. A. Flavahan, Y. Drier, B. B. Liau, S. M. Gillespie, A. S. Venteicher, A. O. Stemmer-Rachamimov, M. L. Suva, and B. E. Bernstein, “Insulator dysfunction and oncogene activation in idh mutant gliomas,” Nature, vol. 529, no. 7584, pp. 110–114, 2016.

[30] J. R. Dixon, I. Jung, S. Selvaraj, Y. Shen, J. E. Antosiewicz-Bourget, A. Y. Lee, Z. Ye, A. Kim, N. Rajagopal, W. Xie, Y. Diao, J. Liang, H. Zhao, V. V. Lobanenkov, J. R. Ecker, J. A. Thomson, and B. Ren, “Chromatin architecture reorganization during stem cell differentiation,” Nature, vol. 518, pp. 331–336, feb 2015.

[31] G. Fudenberg, M. Imakaev, C. Lu, A. Goloborodko, N. Abdennur, and L. A. Mirny, “Formation of Chromosomal Domains by Loop Extrusion,” Cell Reports, vol. 15, pp. 2038–2049, may 2016.

[32] A. L. Sanborn, S. S. P. Rao, S.-C. Huang, N. C. Durand, M. H. Huntley, A. I. Jewett, I. D. Bochkov, D. Chinnappan, A. Cutkosky, J. Li, K. P. Geeting, A. Gnirke, A. Melnikov, I. D. McKenna, E. K. Stamenova, E. S. Lander, and E. L. Aiden, “Chromatin extrusion explains key features of loop and domain formation in wild-type and engineered genomes,” Proceedings of the National Academy of Sciences, vol. 112, pp. E6456–E6465, nov 2015.

[33] M. J. Rowley and V. G. Corces, “Organizational principles of 3D genome architecture,” Nature Reviews Genetics, vol. 19, no. 12, pp. 789–800, 2018.

[34] Y. Guo, Q. Xu, D. Canzio, J. Shou, J. Li, D. Gorkin, I. Jung, H. Wu, Y. Zhai, Y. Tang, Y. Lu, Y. Wu, Z. Jia, W. Li, M. Zhang, B. Ren, A. Krainer, T. Maniatis, and Q. Wu, “CRISPR Inversion of CTCF Sites Alters Genome Topology and Enhancer/Promoter Function,” Cell, vol. 162, pp. 900–910, aug 2015.

[35] W. Schwarzer, N. Abdennur, A. Goloborodko, A. Pekowska, G. Fudenberg, Y. Loe-Mie, N. A. Fonseca, W. Huber, C. H. Haering, L. Mirny, and F. Spitz, “Two independent modes of chromatin organization revealed by cohesin removal,” Nature, vol. 551, pp. 51–56, nov 2017.

[36] S. S. P. Rao, S.-C. Huang, B. Glenn St Hilaire, J. M. Engreitz, E. M. Perez, K.-R. Kieffer-Kwon, A. L. Sanborn, S. E. Johnstone, G. D. Bascom, I. D. Bochkov, X. Huang, M. S. Shamim, J. Shin, D. Turner, Z. Ye, A. D. Omer, J. T. Robinson, T. Schlick, B. E. Bernstein, R. Casellas, E. S. Lander, and E. L. Aiden, “Cohesin Loss Eliminates All Loop Domains.,” Cell, vol. 171, pp. 305–320.e24, oct 2017.

[37] G. Fudenberg, N. Abdennur, M. Imakaev, A. Goloborodko, and L. A. Mirny, “Emerging Evidence of Chromosome Folding by Loop Extrusion,” Cold Spring Harbor Symposia on Quantitative Biology, vol. 82, pp. 45–55, 2017.

[38] W. Gan, J. Luo, Y. Z. Li, J. L. Guo, M. Zhu, and M. L. Li, “A computational method to predict topologically associating domain boundaries combining histone marks and sequence information,” BMC genomics, vol. 20, no. 13, pp. 1–12, 2019.

[39] S. Hong and D. Kim, “Computational characterization of chromatin domain boundary-associated genomic elements,” Nucleic acids research, vol. 45, no. 18, pp. 10403–10414, 2017.

[40] E. Sefer and C. Kingsford, “Semi-nonparametric modeling of topological domain formation from epigenetic data,” Algorithms for Molecular Biology, vol. 14, no. 1, p. 4, 2019.

[41] J. Huang, E. Marco, L. Pinello, and G. C. Yuan, “Predicting chromatin organization using histone marks,” Genome Biology, vol. 16, no. 1, pp. 1–11, 2015.

[42] Y. LeCun, Y. Bengio, and G. Hinton, “Deep learning,” nature, vol. 521, no. 7553, pp. 436–444, 2015.

[43] B. Alipanahi, A. Delong, M. T. Weirauch, and B. J. Frey, “Predicting the sequence specificities of dna-and rna-binding proteins by deep learning,” Nature biotechnology, vol. 33, no. 8, pp. 831–838, 2015.

[44] E. Moen, D. Bannon, T. Kudo, W. Graf, M. Covert, and D. Van Valen, “Deep learning for cellular image analysis,” Nature methods, pp. 1–14, 2019.

[45] A. W. Senior, R. Evans, J. Jumper, J. Kirkpatrick, L. Sifre, T. Green, C. Qin, A. Zídek, A. W. Nelson, A. Bridgland, et al., “Improved protein structure prediction using potentials from deep learning,” Nature, pp. 1–5, 2020.

[46] A. Esteva, B. Kuprel, R. A. Novoa, J. Ko, S. M. Swetter, H. M. Blau, and S. Thrun, “Dermatologist-level classification of skin cancer with deep neural networks,” Nature, vol. 542, no. 7639, pp. 115–118, 2017.

[47] V. Gulshan, L. Peng, M. Coram, M. C. Stumpe, D. Wu, A. Narayanaswamy, S. Venugopalan, K. Widner, T. Madams, J. Cuadros, et al., “Development and validation of a deep learning algorithm for detection of diabetic retinopathy in retinal fundus photographs,” Jama, vol. 316, no. 22, pp. 2402–2410, 2016.

[48] M. Komorowski, L. A. Celi, O. Badawi, A. C. Gordon, and A. A. Faisal, “The artificial intelligence clinician learns optimal treatment strategies for sepsis in intensive care,” Nature medicine, vol. 24, no. 11, pp. 1716–1720, 2018.

[49] R. E. Consortium, “Integrative analysis of 111 reference human epigenomes,” Nature, vol. 518, p. 317, feb 2015.

[50] H. Li and R. Durbin, “Fast and accurate short read alignment with burrows–wheeler transform,” bioinformatics, vol. 25, no. 14, pp. 1754–1760, 2009.

[51] M. Martin, “Cutadapt removes adapter sequences from high-throughput sequencing reads,” EMBnet. journal, vol. 17, no. 1, pp. 10–12, 2011.

[52] E. P. Consortium et al., “An integrated encyclopedia of dna elements in the human genome,” Nature, vol. 489, no. 7414, pp. 57–74, 2012.

[53] T. S. Carroll, Z. Liang, R. Salama, R. Stark, and I. de Santiago, “Impact of artifact removal on chip quality metrics in chip-seq and chip-exo data,” Frontiers in genetics, vol. 5, p. 75, 2014.

[54] Y. Zhang, T. Liu, C. A. Meyer, J. Eeckhoute, D. S. Johnson, B. E. Bernstein, C. Nusbaum, R. M. Myers, M. Brown, W. Li, et al., “Model-based analysis of chip-seq (macs),” Genome biology, vol. 9, no. 9, p. R137, 2008.

[55] B. Alberts, Molecular Biology of the Cell. Garland Science, 6th. ed., 2017.

[56] C. M. Bishop and C. M., Neural networks for pattern recognition. Clarendon Press, 1995.

[57] N. Servant, N. Varoquaux, B. R. Lajoie, E. Viara, C.-J. Chen, J.-P. Vert, E. Heard, J. Dekker, and E. Barillot, “HiC-Pro: an optimized and flexible pipeline for Hi-C data processing,” Genome Biology, vol. 16, no. 1, p. 259, 2015.

[58] B. Langmead and S. L. Salzberg, “Fast gapped-read alignment with bowtie 2,” Nature methods, vol. 9, no. 4, p. 357, 2012.

[59] M. Imakaev, G. Fudenberg, R. P. McCord, N. Naumova, A. Goloborodko, B. R. Lajoie, J. Dekker, and L. A. Mirny, “Iterative correction of Hi-C data reveals hallmarks of chromosome organization,” Nature Methods, vol. 9, no. 10, pp. 999–1003, 2012.

[60] F. Ay, T. L. Bailey, and W. S. Noble, “Statistical confidence estimation for Hi-C data reveals regulatory chromatin contacts,” Genome Research, vol. 24, no. 6, pp. 999–1011, 2014.

[61] F. Serra, D. Bau, M. Goodstadt, D. Castillo, G. Filion, and M. A. Marti-Renom, “Automatic analysis and 3D-modelling of Hi-C data using TADbit reveals structural features of the fly chromatin colors,” PLoS Computational Biology, vol. 13, no. 7, pp. 1–17, 2017.

[62] F. L. Dily, D. Bau, A. Pohl, G. P. Vicent, F. Serra, D. Soronellas, G. Castellano, R. H. Wright, C. Ballare, G. Filion, M. A. Marti-Renom, and M. Beato, “Distinct structural transitions of chromatin topological domains correlate with coordinated hormone-induced gene regulation,” Genes & Development, vol. 28, pp. 2151–2162, oct 2014.

[63] C. Y. McLean, D. Bristor, M. Hiller, S. L. Clarke, B. T. Schaar, C. B. Lowe, A. M. Wenger, and G. Bejerano, “GREAT improves functional interpretation of cis-regulatory regions,” Nature Biotechnology, vol. 28, pp. 495–501, may 2010.

[64] Z. Gu, R. Eils, and M. Schlesner, “Complex heatmaps reveal patterns and correlations in multidimensional genomic data,” Bioinformatics, vol. 32, pp. 2847–2849, 05 2016.

[65] A. R. Quinlan and I. M. Hall, “Bedtools: a flexible suite of utilities for comparing genomic features,” Bioinformatics, vol. 26, no. 6, pp. 841–842, 2010.

[66] L. J. Zhu, C. Gazin, N. D. Lawson, H. Pages, S. M. Lin, D. S. Lapointe, and M. R. Green, “Chippeakanno: a bioconductor package to annotate chip-seq and chip-chip data,” BMC bioinformatics, vol. 11, no. 1, p. 237, 2010.

[67] H. Mi, A. Muruganujan, X. Huang, D. Ebert, C. Mills, X. Guo, and P. D. Thomas, “Protocol update for large-scale genome and gene function analysis with the panther classification system (v. 14.0),” Nature protocols, vol. 14, no. 3, pp. 703–721, 2019.

[68] J. Bergstra, D. Yamins, and D. D. Cox, “Making a science of model search: Hyperparameter optimization in hundreds of dimensions for vision architectures,” Jmlr, 2013.

[69] T. Chen and C. Guestrin, “Xgboost,” Proceedings of the 22nd ACM SIGKDD International Conference on Knowledge Discovery and Data Mining - KDD ‘16, 2016.

[70] F. Pedregosa, G. Varoquaux, A. Gramfort, B. Michel, V.and Thirion, O. Grisel, M. Blondel, P. Prettenhofer, R. Weiss, V. Dubourg, J. Vanderplas, A. Passos, D. Cournapeau, M. Brucher, M. Perrot, and E. Duchesnay, “Scikit-learn: Machine learning in Python,” Journal of Machine Learning Research, vol. 12, pp. 2825–2830, 2011.

[71] D. P. Kingma and J. Ba, “Adam: A method for stochastic optimization,” arXiv preprint 1412.6980, 2014.

[72] T. Gao and J. Qian, “Enhanceratlas 2.0: an updated resource with enhancer annotation in 586 tissue/cell types across nine species,” Nucleic acids research, vol. 48, no. D1, pp. D58–D64, 2020.

[73] M. Lizio, J. Harshbarger, H. Shimoji, J. Severin, T. Kasukawa, S. Sahin, I. Abugessaisa, S. Fukuda, F. Hori, S. Ishikawa-Kato, et al., “Gateways to the fantom5 promoter level mammalian expression atlas,” Genome biology, vol. 16, no. 1, p. 22, 2015.

[74] S. Fishilevich, R. Nudel, N. Rappaport, R. Hadar, I. Plaschkes, T. Iny Stein, N. Rosen, A. Kohn, M. Twik, M. Safran, et al., “Genehancer: genome-wide integration of enhancers and target genes in genecards,” Database, vol. 2017, 2017.

[75] Y. Jiang, F. Qian, X. Bai, Y. Liu, Q. Wang, B. Ai, X. Han, S. Shi, J. Zhang, X. Li, et al., “Sedb: a comprehensive human super-enhancer database,” Nucleic acids research, vol. 47, no. D1, pp. D235–D243, 2019.

[76] A. Khan and X. Zhang, “dbsuper: a database of super-enhancers in mouse and human genome,” Nucleic acids research, vol. 44, no. D1, pp. D164–D171, 2016.

[77] D. R. Zerbino, S. P. Wilder, N. Johnson, T. Juettemann, and P. R. Flicek, “The ensemble regulatory build,” Genome biology, vol. 16, no. 1, p. 56, 2015.

[78] R. Kumar, G. Nagpal, V. Kumar, S. S. Usmani, P. Agrawal, and G. P. Raghava, “Humcfs: a database of fragile sites in human chromosomes,” BMC genomics, vol. 19, no. 9, p. 985, 2019.

[79] D. Karolchik, A. S. Hinrichs, T. S. Furey, K. M. Roskin, C. W. Sugnet, D. Haussler, and W. J. Kent, “The ucsc table browser data retrieval tool,” Nucleic acids research, vol. 32, no. suppl 1, pp. D493–D496, 2004.

[80] A. Heger, C. Webber, M. Goodson, C. P. Ponting, and G. Lunter, “Gat: a simulation framework for testing the association of genomic intervals,” Bioinformatics, vol. 29, no. 16, pp. 2046–2048, 2013.

[81] Z. Xu, H. Wang, S. Wei, Z. Wang, and G. Ji, “Inhibition of er stress-related ire1*α*/creb/nlrp1 pathway promotes the apoptosis of human chronic myelogenous leukemia cell,” Molecular immunology, vol. 101, pp. 377–385, 2018.

[82] Z. Wang, C. Zang, J. A. Rosenfeld, D. E. Schones, A. Barski, S. Cuddapah, K. Cui, T.-Y. Roh, W. Peng, M. Q. Zhang, et al., “Combinatorial patterns of histone acetylations and methylations in the human genome,” Nature genetics, vol. 40, no. 7, p. 897, 2008.

[83] P. H. L. Krijger and W. de Laat, “Regulation of disease-associated gene expression in the 3D genome,” Nature Reviews Molecular Cell Biology, vol. 17, pp. 771–782, ec 2016.

[84] J. Henderson, V. Ly, S. Olichwier, P. Chainani, Y. Liu, and B. Soibam, “Accurate prediction of boundaries of high resolution topologically associated domains (tads) in fruit flies using deep learning,” Nucleic acids research, vol. 47, no. 13, pp. e78–e78, 2019.

[85] R. Bargaje, M. P. Alam, A. Patowary, M. Sarkar, T. Ali, S. Gupta, M. Garg, M. Singh, R. Purkanti, V. Scaria, et al., “Proximity of h2a. z containing nucleosome to the transcription start site influences gene expression levels in the mammalian liver and brain,” Nucleic acids research, vol. 40, no. 18, pp. 8965–8978, 2012.

[86] J. Dorier and A. Stasiak, “The role of transcription factories-mediated interchromosomal contacts in the organization of nuclear architecture,” Nucleic acids research, vol. 38, no. 21, pp. 7410–7421, 2010.

[87] G. J. Filion, J. G. van Bemmel, U. Braunschweig, W. Talhout, J. Kind, L. D. Ward, W. Brugman, I. J. de Castro, R. M. Kerkhoven, H. J. Bussemaker, et al., “Systematic W. protein location mapping reveals five principal chromatin types in drosophila cells,” Cell, vol. 143, no. 2, pp. 212–224, 2010.

[88] M. Thomas, R. L. White, and R. W. Davis, “Hybridization of rna to double-stranded dna: formation of r-loops,” Proceedings of the National Academy of Sciences, vol. 73, no. 7, pp. 2294–2298, 1976.

[89] F. Aymard, M. Aguirrebengoa, E. Guillou, B. M. Javierre, B. Bugler, C. Arnould, V. Rocher, J. S. Iacovoni, A. Biernacka, M. Skrzypczak, et al., “Genome-wide mapping of long-range contacts unveils clustering of dna double-strand breaks at damaged active genes,” Nature structural & molecular biology, vol. 24, no. 4, p. 353, 2017.

[90] A. Canela, Y. Maman, S. Jung, N. Wong, E. Callen, A. Day, K.-R. Kieffer-Kwon, A. Pekowska, H. Zhang, S. S. Rao, et al., “Genome organization drives chromosome fragility,” Cell, vol. 170, no. 3, pp. 507–521, 2017.

[91] S. M. Lundberg and S.-I. Lee, “A unified approach to interpreting model predictions,” in Advances in neural information processing systems, pp. 4765–4774, 2017.

[92] S. Schwartz, E. Meshorer, and G. Ast, “Chromatin organization marks exon-intron structure,” Nature structural & molecular biology, vol. 16, no. 9, p. 990, 2009.

[93] R. F. Luco, Q. Pan, K. Tominaga, B. J. Blencowe, O. M. Pereira-Smith, and T. Misteli, “Regulation of alternative splicing by histone modifications,” Science, vol. 327, no. 5968, pp. 996–1000, 2010.

[94] M. J. Carrozza, B. Li, L. Florens, T. Suganuma, S. K. Swanson, K. K. Lee, W.-J. Shia, S. Anderson, J. Yates, M. P. Washburn, et al., “Histone h3 methylation by set2 directs deacetylation of coding regions by rpd3s to suppress spurious intragenic transcription,” Cell, vol. 123, no. 4, pp. 581–592, 2005.

[95] E. P. Nora, A. Goloborodko, A.-L. Valton, J. H. Gibcus, A. Uebersohn, N. Abdennur, J. Dekker, L. A. Mirny, and B. G. Bruneau, “Targeted degradation of ctcf decouples local insulation of chromosome domains from genomic compartmentalization,” Cell, vol. 169, no. 5, pp. 930–944, 2017.

[96] B. Bonev, N. M. Cohen, Q. Szabo, L. Fritsch, G. L. Papadopoulos, Y. Lubling, X. Xu, X. Lv, J.-P. Hugnot, A. Tanay, et al., “Multiscale 3d genome rewiring during mouse neural development,” Cell, vol. 171, no. 3, pp. 557–572, 2017.

[97] M. J. Rowley, M. H. Nichols, X. Lyu, M. Ando-Kuri, I. S. M. Rivera, K. Hermetz, P. Wang, Y. Ruan, and V. G. Corces, “Evolutionarily conserved principles predict 3d chromatin organization,” Molecular cell, vol. 67, no. 5, pp. 837–852, 2017.

[98] D. G. Lupíañez, K. Kraft, V. Heinrich, P. Krawitz, F. Brancati, E. Klopocki, D. Horn, H. Kayserili, J. M. Opitz, R. Laxova, et al., “Disruptions of topological chromatin domains cause pathogenic rewiring of gene-enhancer interactions,” Cell, vol. 161, no. 5, pp. 1012–1025, 2015.

[99] A. Despang, R. Schöpflin, M. Franke, S. Ali, I. Jerković, C. Paliou, W.-L. Chan, B. Timmermann, L. Wittler, M. Vingron, et al., “Functional dissection of the sox9-kcnj2 locus identifies nonessential and instructive roles of tad architecture,” Nature genetics, vol. 51, no. 8, pp. 1263–1271, 2019.

